# Recruitment of SERK co-receptors determines signaling specificity within the systemin peptide family

**DOI:** 10.64898/2026.02.13.705698

**Authors:** Xu Wang, Wei Yuan, Nga-Thanh Pham, Lei Wang, Rong Li, Anke Steppuhn, Detlef Weigel, Annick Stintzi, Georg Felix, Andreas Schaller

**Affiliations:** Department of Plant Physiology and Biochemistry, Institute of Biology, University of Hohenheim, 70593 Stuttgart, Germany; Department of Molecular Biology, Max Planck Institute for Biology, 72076 Tübingen, Germany; ZMBP, University of Tübingen, 72076 Tübingen, Germany; Yazhouwan National Laboratory, Sanya 572024, Hainan, China; Department of Molecular Botany, Institute of Biology, University of Hohenheim, Stuttgart 70593, Germany; Cluster of Excellence GreenRobust, University of Hohenheim, 70593 Stuttgart, Germany; Cluster of Excellence GreenRobust, Max Planck Institute for Biology, 72076 Tübingen, Germany

## Abstract

The phytocytokine systemin (SYS1) regulates defense responses of tomato plants against herbivores and necrotrophic pathogens. For more than three decades, systemin signaling was thought to rely solely on SYS1, although the tomato genome encodes the precursors of at least three additional systemin-like peptides (SYS2, SYS3 and SYS4). We show that these recently discovered peptides are processed in a wound-inducible manner and act as genuine wound signals that activate early defense signaling via the receptor SYR1. Although all systemins depend on SYR1 for perception, the new peptides elicit downstream responses distinct from those triggered by SYS1, including sustained ethylene production and extensive transcriptional reprogramming. Signaling specificity is governed by a single C-terminal residue that determines the ability of SYS peptides to recruit SERK co-receptors into the receptor complex. The resulting increase in complex stability underlies the amplified and prolonged signaling responses observed for the new SYS peptides compared to SYS1. Our findings reveal how subtle sequence changes within the systemin peptide family can remodel receptor–co-receptor interactions to generate ligand-specific signaling outputs. This mechanism allows a single receptor to decode related ligands into distinct physiological responses and may represent a broader principle allowing for the diversification of peptide-mediated signaling in plants.

## Introduction

Systemin (SYS), an 18-amino-acid oligopeptide, was the first bioactive signaling peptide identified in plants. It was discovered in extracts of wounded tomato leaves in a search for factors that induce protease inhibitor expression as part of the defense response against herbivory (Pearce et al., 1991). The accumulation of protease inhibitors in injured plants enhances resistance by impairing the function of digestive proteinases of insect and mammalian herbivores (Green and Ryan, 1972). SYS is synthesized as a larger precursor protein, prosystemin (PS), from which it is proteolytically released (McGurl et al., 1992; Beloshistov et al., 2018). Acting as a phytocytokine, systemin amplifies defense responses at the site of wounding, where it promotes the production of jasmonates as long-range signals to induce defense gene expression in distal tissues (Schaller and Ryan, 1996; Ryan and Pearce, 1998; Wasternack et al., 2006; Howe and Schaller, 2008).

The early cellular responses to SYS resemble those typically triggered by microbe-associated molecular patterns (MAMPs), including extracellular alkalization, plasma membrane depolarization, induction of an oxidative burst, and an increase in cytosolic calcium (Felix and Boller, 1995; Schaller and Oecking, 1999; Li et al., 2025; Zhai et al., 2025). These early responses are activated after signal perception by cell surface receptors. SYS receptors 1 and 2 (SYR1 and SYR2) were identified by screening introgression lines (ILs) between SYS-sensitive *S. lycopersicum* and SYS-insensitive *S. pennellii*, and were characterized as high-affinity and low-affinity receptors of the SYS peptide, respectively (Wang et al., 2018). SYS responses are fully restored in ILs genetically complemented with SYR1, indicating that SYR2 is dispensable for SYS-induced defense (Wang et al., 2018). In fact, SYR2 was recently shown to rather act in a negative feedback loop dampening the wound response when SYS levels rise to excessively high levels (Zhou et al., 2025).

For decades, the tomato genome was thought to encode only a single SYS peptide, derived from the *PS* gene on chromosome 5. Unexpectedly, we recently detected a gene cluster on chromosome 4 encoding potential precursors of five additional SYS-like peptides (Wang et al., 2025). While one of these peptides appeared to be inactive (SYS5), three of them resembled SYS in activity (SYS2-4), and one was shown to act as a SYS inhibitor (antiSYS). As a receptor antagonist, antiSYS counterbalances the activity of agonistic systemins, thereby preventing inadvertent activation of SYR1 signaling (Wang et al., 2025). Therefore, systemin signaling in tomato appears to be more complex than previously appreciated. Discovery of the SYS family prompts critical questions about functional redundancy versus diversification of peptide activity, and about the mechanisms that enable highly similar peptides to trigger unique downstream responses.

In the present study, we characterize three newly identified tomato systemins—SYS2, SYS3, and SYS4—alongside the canonical SYS peptide (SYS1, hereafter). We show that the new SYS peptides are processed from their precursors in a wound-inducible manner to activate SYR1-dependent defense signaling. Using time-series transcriptomics, we find that all SYS peptides elicit broad transcriptional reprogramming, yet each produces unique transcriptional outputs. In addition to SYS1-controlled genes, the new SYS peptides regulate the expression of genes associated with ethylene signaling, innate immunity, and cell death regulation. Consequently, the new SYS peptides drive more sustained ethylene production compared to the transient ethylene burst triggered by SYS1. Mechanistically we show that SYS2, SYS3, and SYS4 recruit SERK co-receptors to SYR1 far more effectively than SYS1. Enhanced receptor complex formation could be attributed to a single amino acid change at the peptide C-terminus. The data indicate that ligand-dependent differences in the avidity of SYR1 receptor complex formation can cause qualitatively different responses, rather than merely quantitatively different signaling output.

Taken together, our findings expand the systemin paradigm from a single peptide-receptor pair to a small peptide family with differentiated signaling outputs. This diversification provides a mechanism by which plants can fine-tune defense intensity and duration, linking wound perception to broader plant innate immunity.

## Results

### A gene cluster encoding three novel systemin peptides

Resembling SYS1 and its precursor PS1, SYS2, SYS3, and SYS4 peptides (SYS2/3/4) appear to be synthesized as prosystemins (PS2/3/4), from which they are proteolytically released. PS2/3/4 are more closely related to each other than to PS1 (Wang et al., 2025) (Suppl. Figure S1A,B). Like PS1, the encoded precursors lack N-terminal secretion signals and are rich in polar or charged residues. The precursors are quite variable in sequence and length, except for the SYS-like peptides at or very close to their C-termini (Figure 1A, Suppl. Figure S1B). The three residues particularly important for SYS1 activity (Ala1, Pro13 and Thr17; (Pearce et al., 1993)) are conserved in SYS2/3/4 suggesting shared functionality. In contrast, the receptor antagonist antiSYS is two amino acids shorter and is thus lacking the essential Thr17, while the inactive SYS5 peptide lacks three of the residues conserved in SYS1-4, including Ala in position one (Figure 1A).

**Figure 1.**
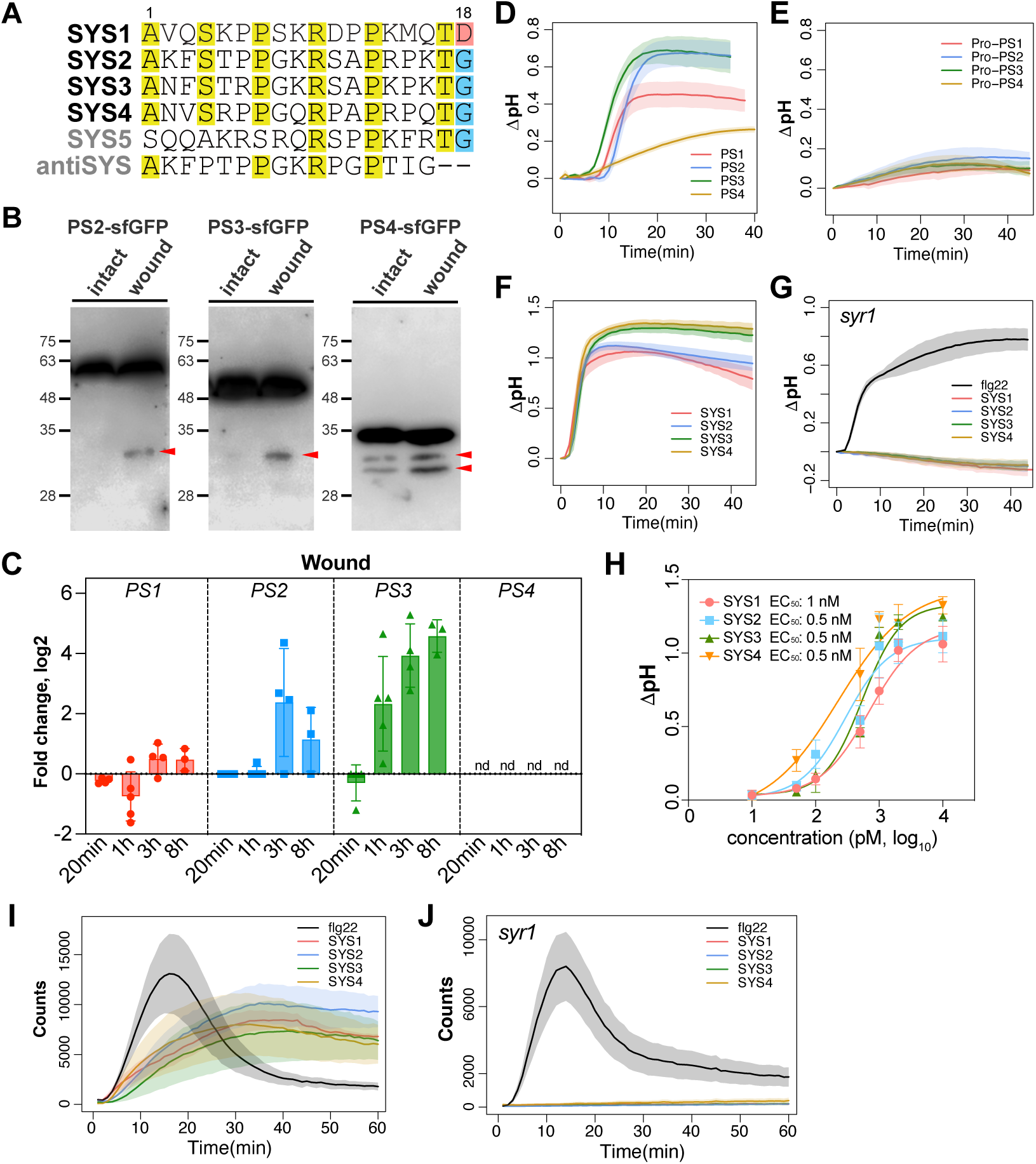
Novel tomato systemin peptides are wound inducible and activate early defense responses in a SYR1-dependent manner. **A**. Sequence alignment of tomato SYS family peptides highlighting conserved residues in yellow and the C-terminal Asp and Gly residues in green and blue, respectively **B.** Wound-induced processing of prosystemin precursors. Western blot analysis of sfGFP-tagged PS proteins in extracts of transgenic (35S::PS-sfGFP) tomato leaves before and 30 min after mechanical wounding. Cleavage products are indicated by red arrows. **C.** Induction of PS gene expression by mechanical wounding. Transcript levels are shown as log2 fold change (log2FC) of normalized transcripts per million (TPM) derived from RNA-seq data. nd, not detected. **D–G**. Medium alkalinization response of suspension-cultured cells to treatment with recombinant PS precursors (**D**), truncated PS precursors lacking the C-terminal SYS region (Pro-PS; **E**), or synthetic SYS peptides (**F,G**). Wild-type and *syr1* mutant cell cultures were used in panels **D-F** and **G**, respectively. Bacterial flg22 was used as a positive control in the *syr1* mutant (**G**). **H**. Dose-response curves and EC_50_ (half-maximal effective concentration) values for SYS1/2/3/4 peptides. I,J. Reactive oxygen species (ROS) burst induced by SYS peptides in leaves of wild-type tomato (I) and *syr1* mutant plants (J). **D-J**. Curves show the mean +/− standard error (SE) of at least 4 independent measurements.

We sought to further characterize the activity of SYS2/3/4, and to investigate their signaling potential compared to SYS1. Since the assumption that SYS2/3/4 peptides are proteolytically released from their precursors is merely based on analogies to SYS1 and antiSYS (Beloshistov et al., 2018; Wang et al., 2025), we first aimed to confirm that the respective precursors are actually processed *in planta*. We therefore expressed sfGFP-tagged PS2/3/4 under control of the CaMV 35S promoter in transgenic tomato lines. Full-length fusion proteins were detected for all PS-sfGFP fusions in unchallenged plants. Precursor processing was observed for PS2-sfGFP and PS3-sfGFP in wounded leaves (Figure 1B). For PS4-sfGFP, processing products increased in abundance after wounding compared to intact leaves. Consistent with their localization at the very C-terminus, only one processing event was observed for the precursors of SYS2 and SYS3. Since SYS4 is not located at the C-terminal end of its precursor (Suppl. Figure 1B), two proteolytic cleavages would be required for peptide release. Consistently, two processed forms were detected for PS4-sfGFP after wounding (Figure 1B). Wound-induced processing of the precursors supports a role for PS2/3/4 as active components of the tomato wound response pathway. Also at the level of gene expression, two of the three new prosystemins were found to be upregulated in response to wounding, as previously reported for *PS1* (McGurl et al., 1992). While overall expression levels did not reach those of *PS1* (Suppl. Fig. 1C), wound-inducibility expressed as fold-induction over the unwounded control was particularly high for *PS2* and *PS3*, starting at 3 and 1 h after wounding, respectively (Figure 1C).

### SYS2/3/4 induce early defense responses in a SYR1-dependent manner

To assess their signaling capacity, we first analyzed the alkalization response of suspension-cultured tomato (*S. peruvianum*) cells to the SYS peptides and their precursors. Recombinant His-tagged PS proteins triggered alkalization of the growth medium starting between six (PS3) and ten (PS2) min after addition to the cell culture (Figure 1D). This activity was lost in PS1/2/3/4 C-terminal deletion constructs indicating that the prodomain is not recognized by cultured tomato cells, and that activity resides in the region harboring the SYS peptides (Figure 1E). Indeed, the synthetic SYS1/2/3/4 peptides all triggered an alkalization response (Figure 1F) which was more rapid and showed a steeper slope of alkalization compared to the respective precursors (compare Figures 1D and 1F). The lag in the response to the precursors indicates that they are not active *per se*, but need to be activated after addition to the culture medium, consistent with proteolytic processing being required for the release of the bioactive peptides. Similar delayed activity due to proteolytic activation in cell culture or leaf extracts has previously been observed for the antagonistic antiSYS peptide (Wang et al., 2025).

Dose-response analyses of the synthetic systemin peptides revealed that the EC_50_ (half maximal effective concentration) was slightly lower for SYS2/3/4 (0.5 nM) compared to SYS1 (1 nM), suggesting that the sensitivity of perception may be somewhat higher for SYS2/3/4 than for SYS1 (Figure 1H). In tomato leaves, all four peptides triggered similar ROS bursts that were slower but more sustained compared to flg22-induced ROS production (Figure 1I). In SYR1-deficient cell cultures and tomato plants generated by CRISPR/Cas9 mutagenesis ((Li et al., 2025) and Suppl. Figure S2), the SYS1/2/3/4-induced alkalization response and ROS production were abolished, while responses to flg22 remained unaffected (Figure 1G, 1J). Conversely, in a gain-of-function experiment using *Nicotiana benthamiana* plants expressing *35S:SYR1-GFP* to confer systemin sensitivity that is lacking in this species, all four SYS peptides elicited ROS production, with higher induction for SYS2 compared to the other SYS peptides (Suppl. Figure S3). These results demonstrated that SYS2, SYS3, and SYS4 are *bona fide* signaling molecules triggering SYS1-like early defense signaling through the SYR1 receptor.

### Herbivore defense is activated by SYS2

To assess their capacity for activating herbivore defense, we compared performance of *Spodoptera exigua* larvae feeding on leaves pre-treated with SYS1 or SYS2 with mock-treated controls. *S. exigua* (the beet armyworm) is a polyphagous herbivore commonly used in plant defense assays. Pre-treatment with either one of the two peptides caused a significant reduction in larval weight gain compared to control leaves (Figure 2A,B). While both peptides had a similar effect on larval growth, mortality early in development was substantially higher in the SYS2 treatment group (Figure 2C, days 4 and 6). The data confirmed that herbivore defenses are activated by both peptides, with a somewhat stronger response for SYS2.

**Figure 2.**
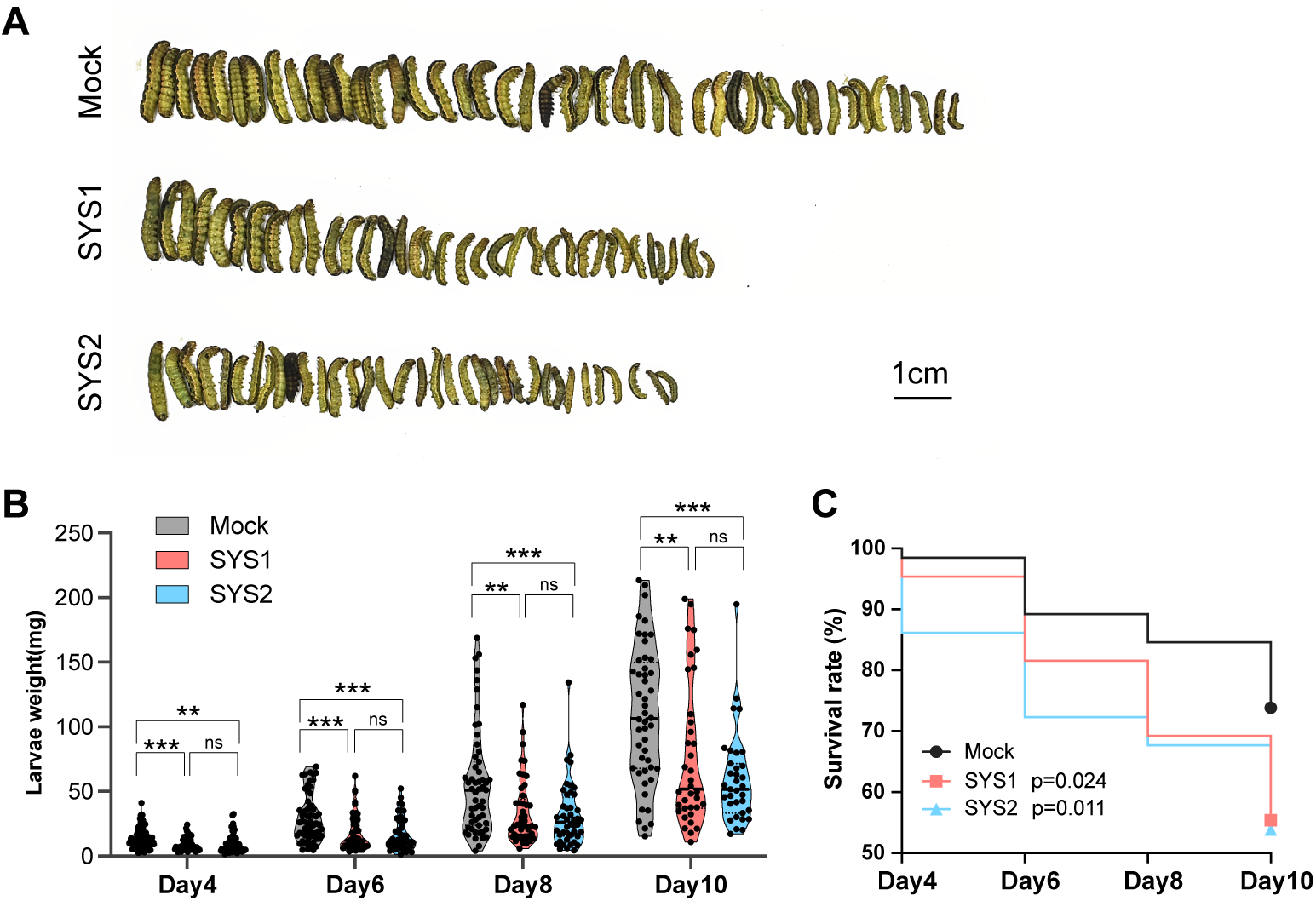
New systemin peptides enhance herbivore resistance against insect larvae. **A**. Representative images of *Spodoptera exigua* larvae after 10 days of feeding on tomato leaves pre-treated with SYS1, SYS2, or mock solution. **B.** Larval fresh weight measured at days 4, 6, 8, and 10 post-feeding. Data are presented as violin plots. Two-tailed Student’s t-test was used to compare mean values for initially 60 larvae; ** and *** indicate significant differences at P < 0.01 and P < 0.001, respectively; ns, not significant. **C**. Larval mortality is higher on SYS2-compared to SYS1- and mock-treated leaves. For each day and treatment, the percentage of surviving larvae is indicated. The Mantel-Cox log-rank test was used to compare overall survival distributions. Significant differences to the mock treatment are indicated by the respective p values.

### Transcriptomic responses differ for different SYS peptides

While the three novel SYS peptides resemble SYS1 in activating SYR1-dependent defense signaling, differences in response magnitude and dose-response kinetics suggested potential for differences in signaling output, including downstream gene expression changes. To investigate this, we conducted a comprehensive transcriptomic analysis by single-end RNA-seq on leaf tissue collected at 0 and 20 minutes, 1, 3, and 8 h after spraying of intact tomato seedlings with the individual SYS peptides (Suppl. Figure S4A,B).

Principal component analysis (PCA) showed that time of treatment was a major source of variance (PC2, 4.9%), while PC3 indicated that each peptide elicited a unique transcriptomic signature (Figure 3A). The number of differentially expressed genes (DEGs) peaked at 1 h for SYS1, SYS3, and SYS4, and at 3 h for SYS2, with most DEGs being up-rather than down-regulated (Figure 3B). The rapid rise and subsequent decline in DEG number over time confirmed that we captured most of the transcriptional response induced by SYS signaling. Comparing transcriptional responses of the four SYS peptides, we observed a substantial core of co-regulated genes at 1 h (40% of all DEGs), and increasingly divergent responses at later time points (Figure 3C). By 8 h, only 5% of the remaining DEGs were common to all four peptides (Figure 3C). Gene expression changes in response to the new SYS peptides included the DEGs of canonical SYS1 but were much stronger in terms of both DEG number and duration. The difference to SYS1 was most pronounced for SYS2, which uniquely induced the expression of over 600 genes (51% of all DEGs) at 3 h after elicitation (Figure 3C).

**Figure 3.**
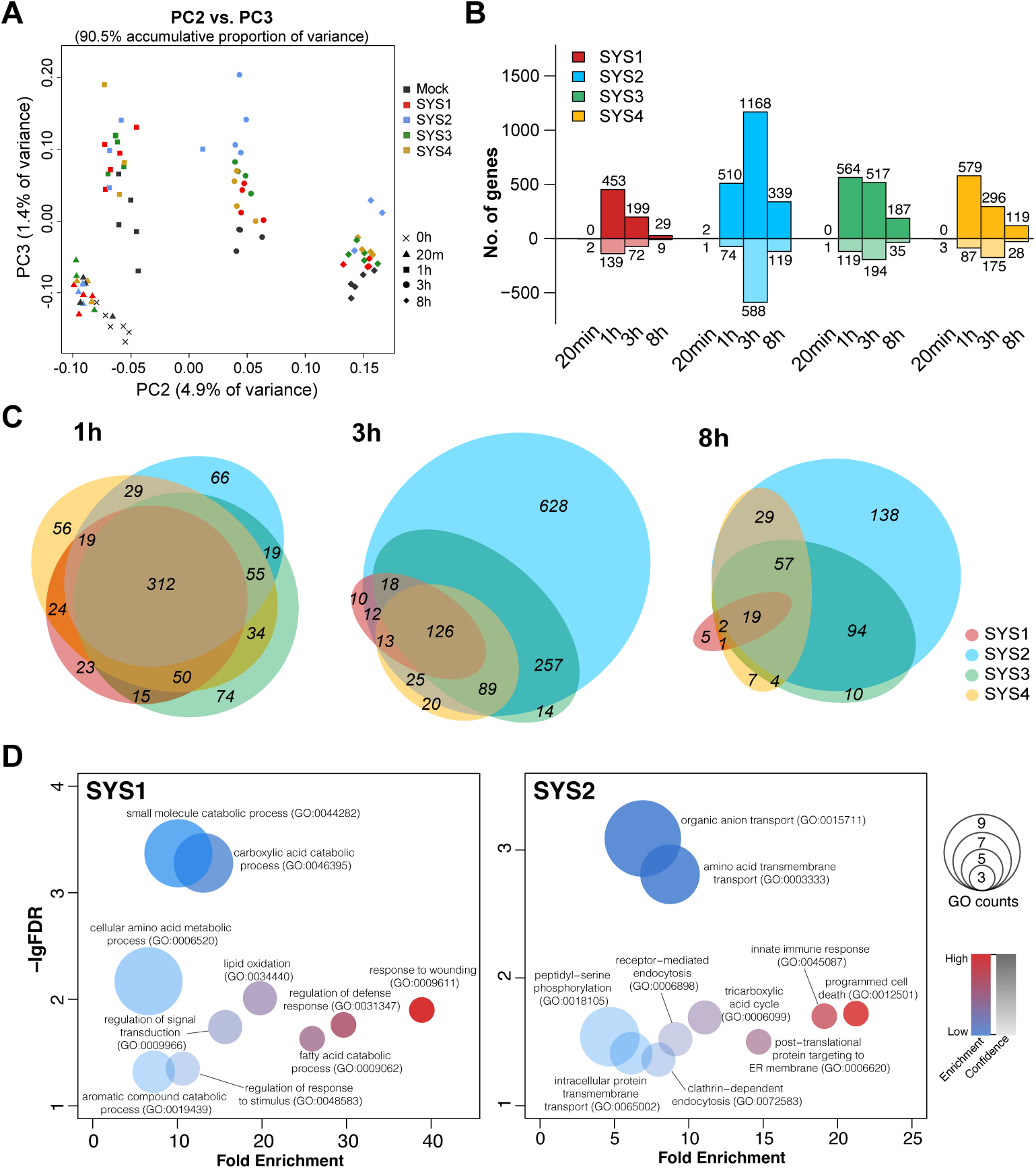
Systemin peptides elicit shared and peptide-specific transcriptional reprogramming. **A.** Principal component analysis (PCA) of RNA-seq data showing global transcriptional dynamics. Variance attributed to ‘time after treatment’ is reflected in PC2, different SYS peptide treatments are separated along PC3. **B.** Number of differentially expressed genes (DEGs) up- and down-regulated for all treatments at different time points. **C.** Euler diagrams illustrating common DEGs, and DEGs specific to the four SYS peptides at 1h, 3hrs and 8hrs after treatment. **D,E**. Bubble plots showing the top 10 GO terms enriched among SYS1-induced (**D**) or SYS2-induced (**E**) DEGs. Enrichment levels and statistical confidence (-lgFDR) are shown on the y- and x-axis, respectively. Bubble size reflects the number of genes associated with the respective GO terms.

While the DEGs shared by SYS1/2/3/4 at the early 1 h time point present a typical wound signaling signature (Supplemental Figure 4C), Gene Ontology (GO) analysis of DEGs detected at 3 h highlighted the contrast between SYS1 and SYS2/3/4 activity. The most highly enriched GO terms for SYS1 were ‘response to wounding’ (39 fold), ‘fatty acid beta-oxidation’ (30 fold), and ‘regulation of defense response’ (30 fold), aligning with its role in jasmonate-dependent wound response signaling (Figure 3D, Supplementary Table S1). In contrast, the top enriched GO terms for SYS2-specific DEGs were ‘programmed cell death’ (21 fold), ‘innate immune response’ (19 fold), and ‘protein targeting to ER membrane’ (15 fold) (Figure 3D; Supplementary Table S2). The novel SYS peptides, particularly SYS2, thus seem to play a broader role in immunity.

To model the dynamic expression patterns across all treatments, we performed linear mixed-model spline (LMMS) fitting followed by k-means clustering (Straube et al., 2015), grouping 15,973 genes into 99 clusters. While many clusters showed shared responses, several exhibited peptide-specific patterns including, for example, cluster 8 (105 genes) and cluster 50 (63 genes). The genes in these two clusters were up- or down-regulated at 3h specifically after SYS2 treatment (Suppl. Figure S4C).

To validate peptide-specific responses, we selected five genes from these clusters for qRT-PCR analysis in an independent experiment: genes for a phosphoserine aminotransferase (Solyc02g082830), a methyltransferase (Solyc04g040180), a NAD(P)-binding protein (Solyc07g047800), ACC oxidase 1 (ACO1, Solyc07g049530), and a sugar transporter (Solyc09g075820). All five genes were strongly induced by SYS2, but not by SYS1, at 3 h (Figure 4A). In contrast, the expression of *Proteinase Inhibitor II* (*PI-II*) as a marker of the canonical jasmonate (JA)-dependent wound response was induced equally by both peptides (Figure 4A). These results confirmed our RNA-seq data, showing that different SYS ligands trigger distinct gene expression responses.

**Figure 4.**
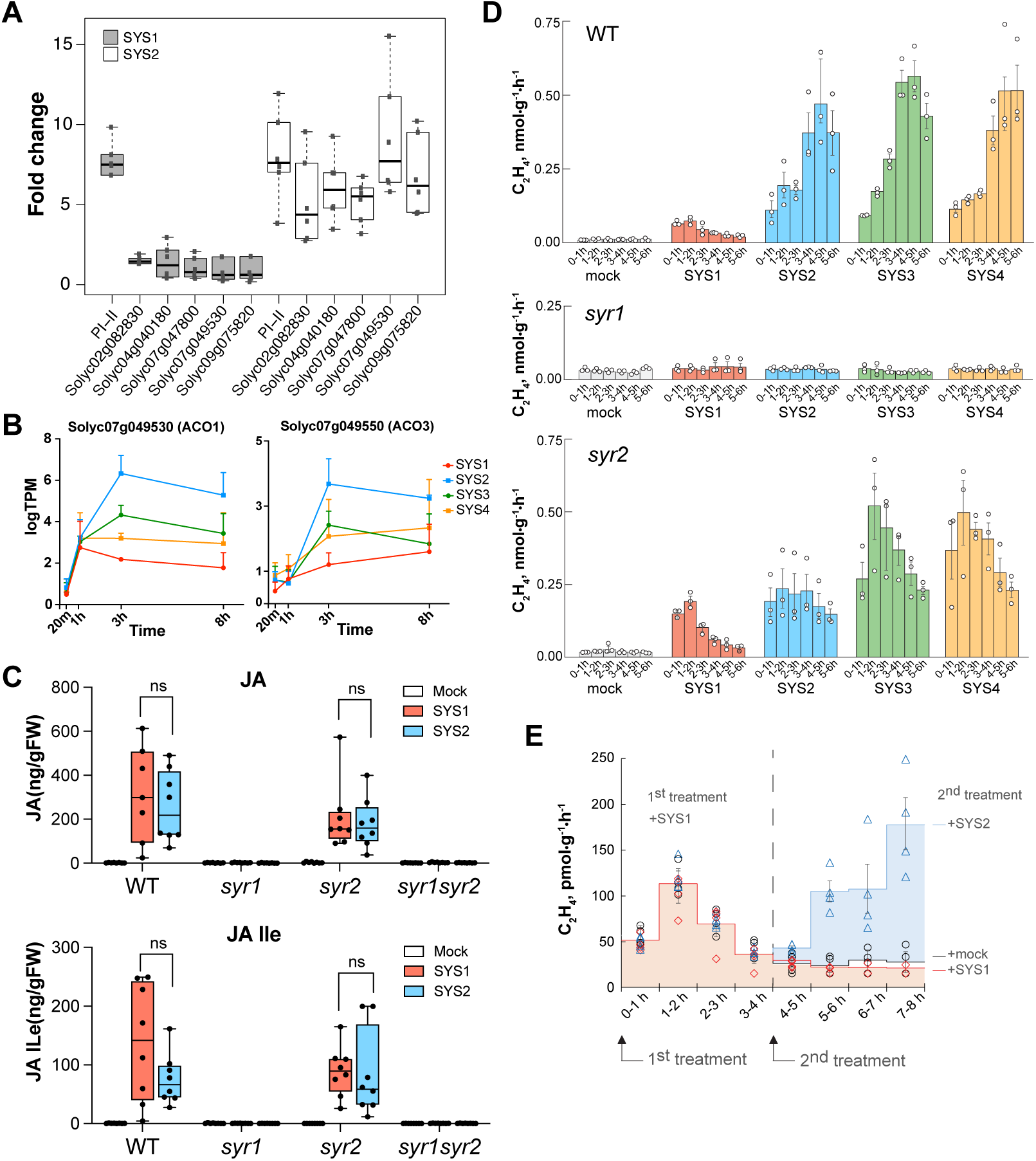
Systemin peptides differentially regulate phytohormone signaling and ethylene biosynthesis. **A.** Transcript levels of five SYS2-specific genes and the *PI-II* wound response marker gene analyzed by quantitative RT-PCR after SYS1 or SYS2 treatment. **B.** Temporal expression profiles of ethylene biosynthesis genes *ACO1* and *ACO3* after peptide treatment. **C.** Levels of JA (top) and JA-Ile (bottom) after 1 hour of SYS1, SYS2 or mock treatment in WT, *syr1*, *syr2* or *syr1syr2* leaves of n = 8 plants; Box plots range from the 25th to 75th percentile with the splitting line at the median and whiskers extending to the minimum and maximum values. ns, non-significant (P ≥ 0.05; two-tailed Student’s t-test) The experiment was performed three times independently with equivalent results. **D, E.** Ethylene emission measured in hourly intervals after treatment with SYS peptides compared to mock in WT, *syr1* or *syr2* mutant lines. (**E**) All samples received a first dose of SYS1 at t = 0, followed by a second treatment with SYS1 (red diamonds), SYS2 (blue triangles), or mock (black circles) at t = 4 hrs.

### SYS2/3/4 elicit sustained ethylene production

SYS2-specific induction kinetics of *ACO1* suggested that the more extensive transcriptional response to SYS2/3/4 compared to SYS1 may involve ethylene synthesis and signaling. Indeed, genes for ethylene synthesis and signaling were induced to higher levels after SYS2/3/4 treatment compared to SYS1 (Figure 4B, Suppl. Figure S5), suggesting peptide-specific modulation of the ethylene pathway. Addressing the involvement of defense hormones in SYS1 vs. SYS2/3/4 signaling, we analyzed ethylene, JA and salicylate (SA) levels in tomato plants after peptide treatment. While no significant increase in SA was observed for any of the peptide treatments (Suppl. Figure S6), JA and JA-Ile were induced to similar levels by SYS1 and SYS2 (Figure 4C). Jasmonates are well established as signal molecules in systemin-mediated wound signaling (Farmer and Ryan, 1992; Li et al., 2002) and their accumulation after both SYS1 and SYS2 treatment is consistent with the observation that core herbivore defense genes including the *PI-II* marker are commonly induced by all systemins (Figures 3C, 4A).

SYS peptides are also known to induce production of the stress hormone ethylene (Felix and Boller, 1995; Wang et al., 2025). However, we observed here that the initial induction of ethylene emission in response to all SYS peptides was followed by a much stronger and longer-lasting increase triggered only by SYS2/3/4 (Figure 4D). This sustained ethylene production is thus a specific feature of SYS2/3/4 signaling and may explain the much larger transcriptomic response to these peptides compared to SYS1, including many ethylene-related genes, particularly at later time points after treatment (Figure 3C, Suppl. Fig. S6)). Interestingly, the induction of jasmonate and ethylene production, like the early responses at the plasma membrane (extracellular alkalization and ROS burst), all relied on SYR1. They were lost in the *syr1* mutant but unaffected in *syr2* (Figs. 4C,D).

### Differences in perception rather than peptide stability account for SYS1 vs SYS2/3/4 signaling specificity

As a first hypothesis potentially explaining the more sustained and stronger response to SYS2/3/4 compared to SYS1, we addressed the possibility that differences in peptide stability accounted for the observed differences in activity. Indeed, SYS1 is known to be unstable in cell culture medium with a half-life of about ∼10 min (Felix and Boller, 1995) due to proteolytic cleavage after Lys14 (Schaller, 1998). While this basic residue (Lys/Arg) in position 14 is conserved in all SYS peptides, only SYS2/3/4 feature Pro in position 15 (Figure 1A). Since most proteases are unable to cleave proline-adjacent bonds (Vanhoof et al., 1995), unspecific degradation may thus be lower for SYS2/3/4, resulting in potentially increased metabolic half-life compared to SYS1.

To test whether peptide instability can explain the comparatively weak and more transient ethylene peak induced by SYS1, we added a second dose of SYS1 four hours after the initial treatment. Interestingly, there was no further ethylene production in response to SYS1 indicating that peptide stability is not the limiting factor (Figure 4E). Surprisingly, SYS1-pretreated cells still responded with increased ethylene production to a second dose of SYS2, 3, or 4 (Figure 4E, Suppl. Figure 7). The observation of cells being refractory to SYS1 after SYS1 pretreatment but fully responsive to SYS2/3/4 revealed that the plant’s perception system distinguishes between SYS1 on the one hand and SYS2/3/4 on the other, and that sustained ethylene production is a specific feature of SYS2/3/4 signaling output, rather than an artifact of peptide persistence.

### SYS peptides differentially recruit SERK co-receptors to SYR1

Our data indicated that SYS1 and SYS2/3/4 perception triggers similar early responses at the plasma membrane (ROS burst, alkalization response), but different signaling outputs in terms of ethylene production and defense gene activation. Since all SYS-induced defense responses are mediated by the same primary receptor, SYR1 (Figure 1G,J; Figure 3D), we hypothesized that ligand-specific differences in co-receptor recruitment or receptor complex stability may account for the different types of responses. To test this, we immuno-precipitated SYR1 complexes from stably transformed 35S:SYR1-GFP tomato plants treated with either SYS1 or SYS2, followed by mass spectrometry (Co-IP/MS). In SYS2-treated samples, we robustly detected somatic embryogenesis receptor kinases (SERKs), which are well-established co-receptors for LRR-RKs including SYR1 (Figure 5A)(Cho et al., 2024; Wang et al., 2025; Zhou et al., 2025). The major isoforms detected after SYS2 treatment were SERK3A and 3B (10 % of all peptides associated with SYR1) with a minor contribution from SERK1 (0.45 % of all peptides). Surprisingly, SERK3A/3B-derived peptides were barely detectable in the immuno-precipitate of SYS1-treated samples (0.02 % of all peptides; Figure 5A).

**Figure 5.**
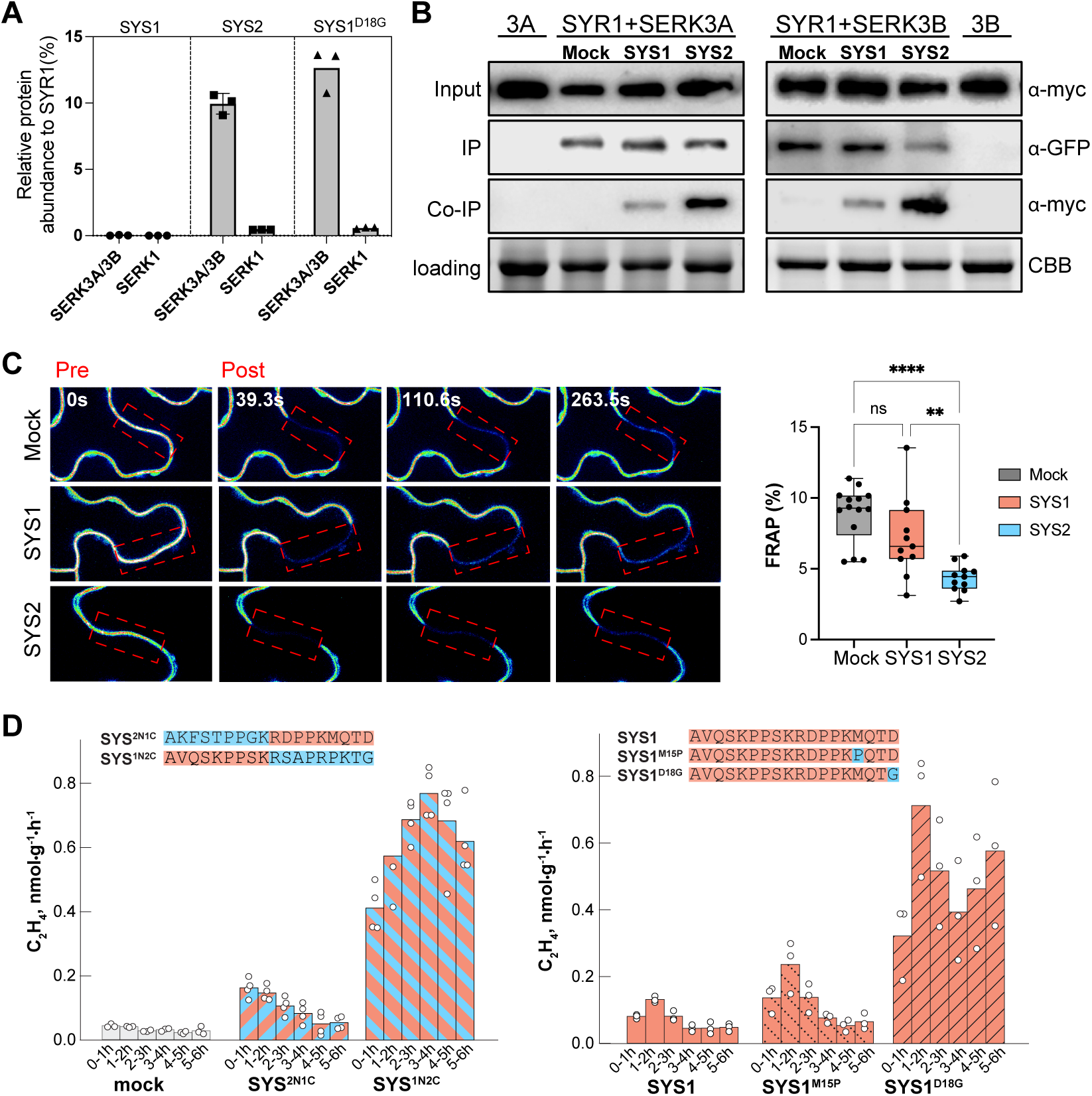
Systemin peptides differentially recruit SERK co-receptors to SYR1. **A.** Co-IP/MS analysis of *35S:SYR1-GFP* overexpression plants treated with SYS1, SYS2 or SYS1^D18G^. SERK proteins (SERK3A/SERK3B and SERK1) were specifically detected 5 min after SYS2 and SYS1^D18G^ treatments. **B.** Co-IP assay of SYR1 interaction with SERK3A or SERK3B upon SYS1 or SYS2 treatment. SYR1 was co-expressed as GFP fusion with Myc-tagged SERK3A or SERK3B in *N. benthamiana* leaves. Total protein extracts from leaves treated with 100 nM SYS1 or SYS2 were immunoprecipitated with GFP-trap beads, then detected with anti-GFP and anti-Myc on western blots. An anti-Myc blot and Coomassie (CBB)-stained gel of the input fractions are shown as loading controls. The assay was performed three times with equivalent results. C. Representative confocal images show color-coded GFP fluorescence intensities before (pre) and after (post) photobleaching. Fluorescence recovery (FRAP) in the photobleached areas (dotted boxes) after 300 seconds is shown in % of the initial fluorescence level. Box plots range from the 25th to 75th percentile with the splitting line at the median. Whiskers extend to the minimum and maximum. Two-tailed Student’s t-test was used for pairwise comparisons of the mean of n = 11 measurements; ** and **** indicate significant differences at P<0.01 or P<0.0001, respectively; ns, non-significant (P ≥ 0.05). **D.** ethylene emission induced by chimeric SYS peptides comprising N- and C-terminal halves of SYS1 and SYS2 as indicated (SYS^2N1C^ and SYS^1N2C^; left), and single-site substituted SYS1 derivatives (SYS1^Q16K^ and SYS1^D18G^; right). The sequences of tested peptides are shown on top, with SYS1 and SYS2-derived residues highlighted in red and blue, respectively.

Suspecting that robust detection of SERK-SYR1 interaction after SYS1 treatment may be limited by the sensitivity of the Co-IP/MS assay, we performed pulldown experiments with *N. benthamiana* leaves transiently co-expressing SYR1-GFP with SERK3A or 3B. Consistent with the Co-IP/MS data, much stronger interaction of SYR1 with SERK3A/3B was observed upon addition of SYS2 compared to SYS1 (Figure 5B). Comparing all four SYS peptides, the ability to recruit SERKs into the receptor complex (Suppl. Figure S8) was found to correlate with the extent of peptide-induced transcriptional reprogramming (Figure 3B,C), being highest for SYS2 (1168 DEGs upregulated at 3 hrs) compared to SYS3 (517 DEGs), SYS4 (296 DEGs) and SYS1 (199 DEGs).

Aiming to confirm apparently ligand-specific differences in receptor complex formation, we performed Fluorescence Recovery After Photobleaching (FRAP) assays in 35S:SYR1-GFP expressing tomato plants. While SYS1 treatment slightly reduced SYR1-GFP mobility (7.2% recovery) compared to the mock control (8.9%), SYS2 treatment immobilized the receptor more strongly (4.3% recovery) (Figure 5C). The results are consistent with the formation of a more stable receptor complex with SYS2 compared to SYS1 as the ligand.

We then asked which specific features of the SYS peptides may account for the observed difference in co-receptor recruitment and, consequently, signaling output. For this we tested chimeric systemins constructed from sequences of SYS1 and SYS2 (Fig. 5D) and found that the ethylene response pattern is determined by the peptides’ C-terminal part. Ethylene emission was transient for the peptide ending with the SYS1 sequence (SYS^2N1C^), and persistent when ending with SYS2 (SYS^1N2C^). Next, we tested whether replacing single residues of SYS1 with their SYS2 counterparts could convert the response pattern from transient to persistent. Two residues in the C-terminal part of SYS1 - Met15 and Asp18 - differ in SYS2/3/4, where these positions are occupied by Pro and Gly, respectively (Figure 1A). While substitution of Met15 with Pro (SYS1^M15P^) had no effect, replacement of the C-terminal Asp by Gly (SYS1^D18G^) recapitulated the sustained ethylene emission response characteristic of SYS2/3/4 (Figure 5E). Strikingly, the D18G substitution also greatly enhanced the ability of SYS1 to recruit SERK3A/B into the receptor complex, as indicated by Co-IP-MS (Figure 5A) and pulldown experiments (Suppl. Figure 8). These results directly link the capacity for co-receptor recruitment to downstream signaling output.

## Discussion

### SYS peptides differ in their downstream signaling potential

The discovery of systemin (SYS1) in 1991 and its precursor prosystemin (PS) in 1992 (Pearce et al., 1991; McGurl et al., 1992) introduced peptides into the suite of plant hormones, establishing a new signaling paradigm in plants. *PS* has always been considered a single-copy gene, but the recent identification of a *PS* gene cluster in tomato fundamentally changed this view (Wang et al., 2025). Our study builds upon this discovery by functionally characterizing the signaling network orchestrated by the SYS peptide family. We confirm that formation of the new SYS peptides (SYS2/3/4) is wound-inducible, consistent with the canonical SYS1 pathway (Figure 1B,C). We found that all SYS peptides signal through SYR1 to trigger similar early cellular immune responses (alkalinization response (Figure 1F,G), ROS burst (Figure 1I,J) and the early phase of ethylene formation (Figure 4D). Interestingly however, SYS signaling diverged thereafter, resulting in peptide-specific differences in ethylene production (Figure 4D,E) and transcriptional reprogramming (Figure 3).

The elevated and more sustained emission of ethylene in response to SYS2/3/4 compared to SYS1 (Figure 4D) is likely facilitated through the specific up-regulation of key ethylene-biosynthesis and perception genes (Figure 4A,B, Suppl. Figure S6). While some components of the ethylene pathway were uniformly induced by all SYS peptides, including *ACO2* (Solyc12g005940), *ACO6* (Solyc02g036350), and one ACC-synthase (ACS, Solyc03g007070), others exhibited peptide-specific regulation. Notably, *ACO1* (Solyc07g049530) and *ACO3* (Solyc07g049550) were specifically upregulated in response to SYS2/3/4 treatments (Figure 4B). The traditional view holds that ACS rather than ACO is the rate-limiting enzyme in ethylene biosynthesis (Adams and Yang, 1981). However, growing evidence indicates that ACO can become rate-limiting for ethylene production during various developmental and stress responses (Houben and Van de Poel, 2019). Our findings support this emerging paradigm by demonstrating that SYS peptides precisely control the expression of specific *ACO* family members. The peptide-specific induction of *ACO1* and *ACO3* correlates with the sustained ethylene emission observed in SYS2/3/4-treated plants, suggesting that differential regulation of *ACO* gene expression shapes ethylene dynamics in response to different danger signals. Sustained ethylene production in response to SYS2/3/4 correlates with distinct downstream transcriptomic responses related to innate immunity and programmed cell death (PCD).

### Role of SERKs serving as co-receptor in SYS-SYR1 signaling

Our data show that the systemin perception machinery can discriminate between SYS1 and the newly identified SYS peptides (Figure 4E), even though all four peptides signal through a single, shared receptor. Indeed, the responses triggered by the different SYS peptides all relied on SYR1 as the primary receptor (Figures 1G,J and 4C,D). The observed ligand-specific differences with respect to ethylene production (Figures 4D and 5D) and defense gene activation (Figure 3, Suppl. Figure S5) therefore suggest a model where different SYR1 receptor complexes, with distinct signaling capacities, are formed in response to different SYS peptides.

Receptor complexes involving leucine-rich repeat receptor kinases (LRR-RKs) or receptor-like proteins (LRR-RLPs) are formed upon ligand-induced heterodimerization with co-receptors of the SERK family (Han et al., 2014; Macho and Zipfel, 2014). Individual SERK members exert distinct functions with a high degree of functional plasticity and specificity (Ma et al., 2016). Tomato has three SERKs (SERK1, SERK3A, and SERK3B) with defined, often non-redundant functions in immunity. For instance, SERK3A and SERK3B, but not SERK1, serve as co-receptors of the flagellin receptor FLS2 and the cold shock protein receptor CORE for the recognition of bacterial MAMPs (Peng and Kaloshian, 2014; Wang et al., 2016; Cho et al., 2024). While all three tomato SERKs contribute to Ve1-mediated resistance to Verticillium wilt (Fradin et al., 2009; Fradin et al., 2011), only SERK1 and SERK3A are required for the hypersensitive response and resistance to *Passalora fulva* (Postma et al., 2016), and Mi-1-mediated defense against aphids depends only on SERK1 (Mantelin et al., 2011).

Multiple studies have confirmed that SERKs are co-receptors of SYR1. All three tomato SERKs were reported to interact with SYR1 after SYS1 treatment (Cho et al., 2024; Wang et al., 2025; Zhou et al., 2025). Genetic evidence further confirmed involvement of SERK1 and SERK3A, while SERK3B seemed dispensable, for SYR1-dependent immune responses induced by SYS1 (Cho et al., 2024; Zhou et al., 2025). Therefore, formation of a different receptor complex in response to SYS2/3/4, by specific recruitment of SERK3B or of another, unrelated co-receptor, could potentially account for the observed ligand-dependent differences in signaling output. However, our data do not support such a scenario. Using a SYR1-GFP fusion as bait, Co-IP/MS robustly pulled down SERK3A/3B with a minor contribution of SERK1 specifically after SYS2 treatment (Figure 5A). When SYS1 was used as the ligand, neither SERKs nor any other co-receptors were co-precipitated. However, using the ectopic *N. benthamiana* expression system, we were able to detect SYS1-triggered interaction of SYR1 with both, SERK3A and SERK3B. While this interaction was markedly weaker than that induced by SYS2/3/4, it confirms that SERKs are part of the SYS1-induced receptor complex. These results are in line with previous studies reporting SYS1-induced SYR1-SERK interaction (Cho et al., 2024; Wang et al., 2025; Zhou et al., 2025).

While receptor (SYR1) and co-receptor (SERK3A/3B) were found to be the same for all SYS peptides, ligand-specific differences were observed with respect to stability of the receptor complex. Indeed, Co-IP/MS (Figure 5A) and pull-down experiments (Figure 5B, Suppl. Figure S8) indicated that the interaction of SYR1 with SERK3A and 3B is much stronger in presence of SYS2/3/4 compared to SYS1. Consistent with higher stability of the SYS2-compared to the SYS1-induced receptor complex, FRAP assays revealed stronger reduction in mobility of SYR1-GFP in presence of SYS2 than with SYS1 (Figure 5C). We conclude that ligand-dependent differences in complex stability, particularly increased stability of the SYS2-SYR1-SERK3A/3B complex compared to SYS1-SYR1-SERK3A/3B, account for SYS2-specific transcriptional responses and ethylene production. This conclusion is supported by structure predictions (AlphaFold3), resulting in a high-confidence model for the ternary complex comprising SYR1, SERK3A and SYS2 (ipTM = 0.83; Figure 6A). The predicted complex resembles experimentally solved structures of many other LRR-RK-SERK complexes in which the respective peptide ligand (FLS2 (Sun et al., 2013), TDIF (Zhang et al., 2016), IDA (Santiago et al., 2016), MIK2 (Jia et al., 2024; Wu et al., 2024)) is sandwiched between the receptor’s LRR domain and the recruitment loop of the SERK co-receptor. In contrast, no such model was obtained with SYS1 as the ligand (ipTM = 0.31). While SYS1 was predicted to bind, similarly to SYS2, to the inner concave surface of the SYR1 LRR, no interactions were predicted between SYS1 and SERK3A (Figure 6C). Modeling predictions are thus consistent with our conclusion that SYS1 does not facilitate binding of the co-receptor to the same extent as SYS2 does.

**Figure 6.**
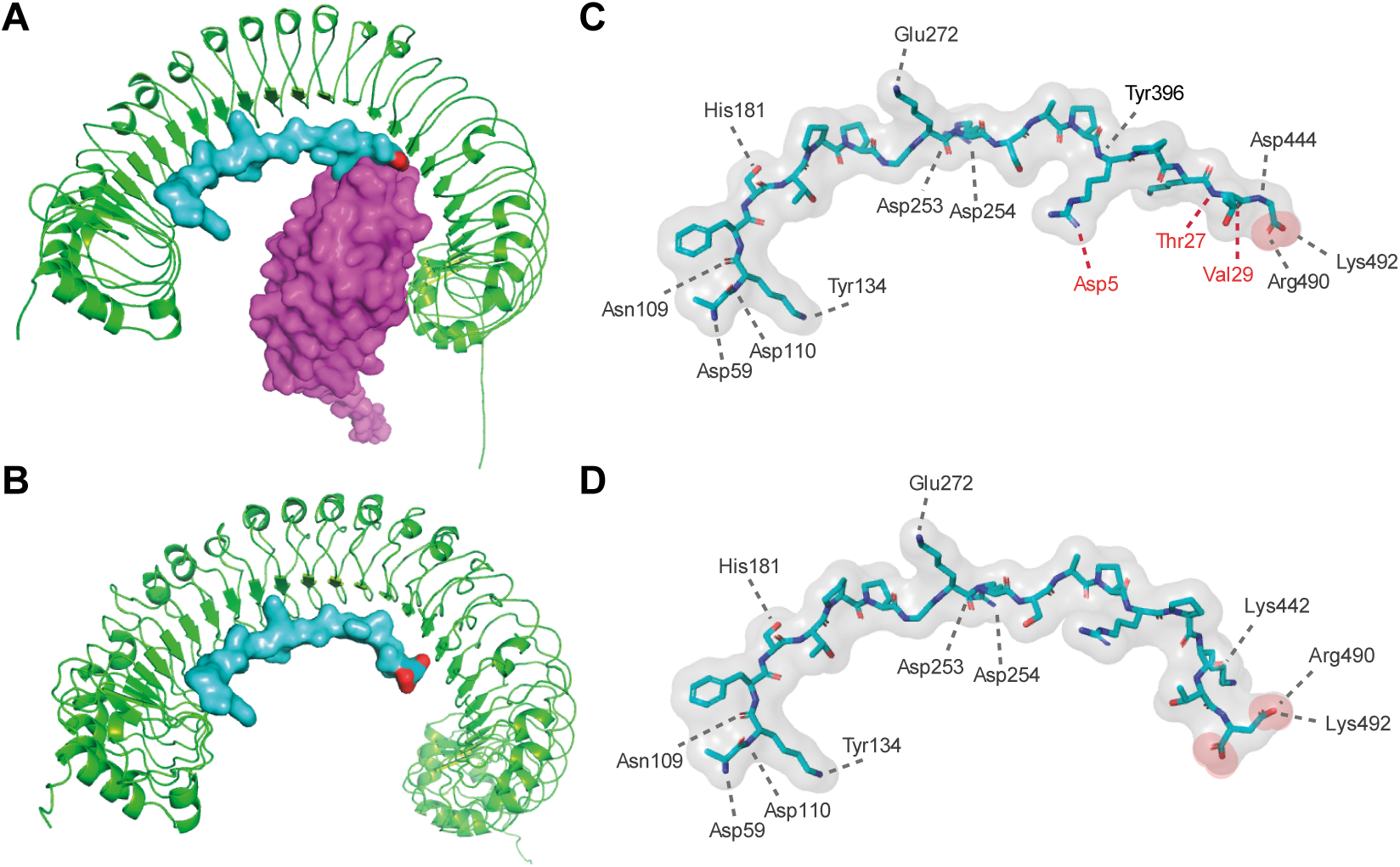
Structural modelling supports a role for systemin’s C-terminus in co-receptor recruitment and receptor complex stability. Structural models (Alphafold3; https://alphafoldserver.com/) of SYS2 (**A**) and SYS2^G18D^ (**B**) as bound in the predicted SYR1-SERK3A complexes (receptor in green, co-receptor in magenta, ligand in blue with carboxyl groups of C-terminal residues highlighted in red). Interface predicted template modeling (ipTM) scores were 0.83 for SYS2-SYR1-SERK3A and 0.56 for SYS2^G18D^-SYR1-SERK3A (modeling scores for SYS2-SYR1-SERK3B and SYS1-induced complexes are compiled in Suppl.Table S4). Consistent with the low confidence score for the complex shown in B, no meaningful interaction with SERK3A was predicted. Since Alphafold3 placed the co-receptor on the opposite convex surface of SYR1 it was omitted here for clarity. **C,D**. Close-up view of the ligands. Peptide backbone and side chains are shown as sticks (carbon in cyan, oxygen in red, nitrogen in blue), the surface in gray, except for the carboxyl groups of the C-terminal Gly (**C**) and Asp (**D**) residues which are highlighted in red). SYR1 residues predicted to form hydrogen bonds or salt bridges to SYS2 (**C**) or SYS2^G18D^ (**D**) are shown in black, those formed with SERK3A in red. Figures were generated in PyMol.

Differences between SYS1- and SYS2/3/4-induced binding of SERKs and receptor complex stability may have important ramifications for the regulation of SYS signaling. Canonical SYS1 signaling was recently shown to be regulated in a negative feed-back loop by SERK availability. When SYS1 accumulates to excessively high levels during a sustained wound response, it binds to the low-affinity SYR2 receptor. Consequently, SERKs are sequestered into the unproductive SYR2-SERK complex and no longer available for signaling through SYR1 (Zhou et al., 2025). Given that SYS2/3/4 bind SERKs more strongly than SYR1, we would predict that SYS2/3/4 signaling is less sensitive to SYR2-mediated inhibition compared to SYS1. The same prediction applies to antiSYS-mediated inhibition of SYS signaling. antiSYS is important in unwounded plants preventing inadvertent activation of defense responses. When SYS levels are low prior to herbivore attack, the antagonistic peptide binds to SYR1 but fails to recruit SERKs into the complex (Wang et al., 2025). This antagonistic activity is overcome when SYS levels rise after herbivory to levels sufficient to outcompete antiSYS from its binding site on SYR1, thereby facilitating receptor complex formation (Wang et al., 2025). With higher stability of the SYS2/3/4 compared to the SYS1-induced complex, we would predict that antiSYS-mediated inhibition is overcome more easily (at lower concentrations) by SYS2/3/4 than by SYS1.

### The SYS C-terminus governs receptor complex formation and signaling specificity

Early biochemical studies indicated that perception and signaling of SYS1 rely on its N-terminal region for binding to a cell surface receptor that had not been identified at that time, and on its C-terminus for receptor activation and signaling (Meindl et al., 1998). This two-step mode of perception follows the ‘address-message’ concept initially proposed for animal neuropeptides (Schwyzer, 1977; Schwyzer, 1987), and later applied in plants to describe the interaction of peptide ligands with cognate LRR-RKs (Meindl et al., 2000; Sun et al., 2013; Macho and Zipfel, 2014). According to this concept, selective binding of the receptor and its activation are mediated by different domains of the ligand, the ‘address’ and ‘message’ domains, respectively. Structure-activity analyses support this concept for SYS1. While C-terminal deletions abolish SYS1 activity (‘the message’), such truncated peptides still bind (via ‘the address’) to the receptor and act as potent receptor antagonists (Pearce et al., 1993; Meindl et al., 1998; Wang et al., 2018).

In addition to the C-terminal Asp^18^, deletion of which renders SYS1 inactive, substitutions of Pro^13^ or Thr^17^ also have a major impact on SYS1 activity (Pearce et al., 1993). While Pro^13^ and Thr^17^ are conserved in the newly identified SYS2/3/4 peptides, glycine instead of aspartate is present in position 18 (Figure 1A). This divergence at the extreme C-terminus suggested that the residue in this position could be particularly important for peptide specificity. Indeed, our functional assays showed that introducing the SYS1^D18G^ mutation was sufficient to produce a SYS2-like phenotype, promoting strong SERK recruitment (Figure 5A, Suppl. Figure S8) and inducing the sustained ethylene burst characteristic of SYS2 signaling (Figure 5E). These results demonstrate that the C-terminal residue governs ligand-specific differences in stability of the SYR1-SERK complex and signaling output.

AlphaFold3 simulations of the SYS2-SYR1-SERK3A complex (Figure 6A,B) provide a plausible explanation for the involvement of SYS2’s C-terminus in receptor complex formation. The predicted model closely resembles experimentally solved structures of several plant peptide-receptor complexes (Song et al., 2017). Similar to the cryo-EM structure of the SCOOP12-MIK2-BAK1 complex (Jia et al., 2024), for example, the model shows the C-terminus of SYS2 sandwiched between the LRR domain of SYR1 and the recruitment loop of SERK3A (residues 50 – 61; Figure 6A). The C-terminal carboxylate of SYS2 forms salt bridges with Arg490 and Lys492 of SYR1 (Figure 6B). These two basic residues separated by valine (RVK) correspond to the RXR motif that is critically important in MIK2 and many other LRR-RKs for binding of their ligand’s C-termini (Song et al., 2017; Jia et al., 2024; Wu et al., 2024). SYS2 also interacts with SERK3A, thereby contributing directly to co-receptor binding. Two hydrogen bonds are predicted between the main chain carbonyl and amide of Thr17^SYS2^ with Thr27 and Val29 in the recruitment loop of SERK3A (Figure 6B). Equivalent main chain interactions between the penultimate residue of the ligand and the recruitment loop of the co-receptor have also been observed e.g. in the TDR-CLE41-SERK2 and HAESA-IDA-SERK1 complex structures (Santiago et al., 2016; Zhang et al., 2016; Song et al., 2017). In the predicted SYS2-SYR1-SERK3A complex, an additional salt bridge is formed between the side chain of Lys14^SYS2^ and Asp5^SERK3A^ (Figure 6B).

Consistent with the experimentally observed relevance of the C-terminal residue for co-receptor recruitment and signaling activity, substituting Gly18 in SYS2 with Asp (the native terminal residue of SYS1) induced an alternative binding mode. In the predicted complex, the Asp18 side chain of SYS2^G18D^ —rather than the backbone carboxylate of Gly18—interacts with the RVK motif of SYR1. Alternative binding of the peptide’s C-terminus causes a conformational change precluding the interaction of SYS2^G18D^ with SERK3A (Figure 6C,D). Importantly, the change in conformation affected only the last five amino acids of the ligand. All hydrogen bonds predicted in the SYS2-SYR1-SERK3A complex between the N-terminal part of SYS2 and SYR1 were also observed for SYS2^G18D^ (Figure 6C,D).

With all N-terminal interactions retained, the ‘address’ would be the same for SYS1 and SYS2, efficiently targeting the SYR1 receptor. Structural modeling suggests that the last amino acid directs the binding mode of the peptide’s C-terminus. The C-terminal conformation of the ligand directly affects co-receptor recruitment and receptor complex stability which, in turn, determine signaling output. Therefore, the ‘message’ sent differs for different SYS peptides. Systemins with Gly compared to Asp in the ultimate position are more effective in SERK recruitment. Stabilization of the receptor complex translates into enhanced immune signaling including more persistent ethylene production and massive transcriptional reprogramming.

This expanded address-message concept explains how a group of related peptides, targeting the same receptor, achieves peptide-specific signaling output. We propose that this concept may not only apply to systemins, but may be more generally applicable explaining differences in activity also in other families of plant signaling peptides.

## Experimental procedures

### Plant material and growth conditions

Tomato (*Solanum lycopersicum* cv. UC82B) and *Nicotiana benthamiana* plants were cultivated in growth chambers at 26 °C under a 16 h photoperiod, 100 µmol m⁻² s⁻¹ light intensity, and 65% relative humidity. *Solanum peruvianum* cell suspension cultures were maintained at 26 °C and 120 rpm by weekly transfer to fresh medium (Wang et al., 2024). Transgenic tomato lines overexpressing 35S:PS-sfGFP were generated via *Agrobacterium*-mediated transformation as described (Li et al., 2025). Briefly, cotyledon segments from 10-day-old etiolated seedlings were co-cultivated for two days with *A. tumefaciens* strain GV3101 carrying the expression constructs in a binary vector with the *nptII* gene for kanamycin resistance as selection marker. Resistant plants were regenerated from calli and the T1 progeny was tested for segregation of the resistance marker and GFP fluorescence. Homozygous T2 lines for each construct were selected for further experiments.

### Molecular cloning

Prosystemin open reading frames (ORFs) were amplified from *S. lycopersicum* leaf cDNA synthesized from total RNA of 3-week-old seedlings using Q5 High-Fidelity DNA polymerase (New England Biolabs) and gene-specific primers (Supplementary Table S3). PCR products were cloned between an N-terminal FLAG tag and a C-terminal sfGFP tag into pART7. The entire expression cassette comprising the CaMV 35S promoter and *OCS* terminator was cut out of pART7 with *NotI* and subsequently ligated into the binary vector pART27 (Gleave, 1992). For bacterial expression, the ORFs of full-length prosystemins or their prodomain regions were cloned into pET-33B+ to generate N-terminally His-tagged proteins. Constructs were transformed into *E. coli* strain BL21 RIL for protein production. All ORFs were verified by Sanger sequencing. Primer sequences are provided in Supplementary Table S3.

### Purification of His-tagged proteins

*E. coli* BL21 RIL cells were grown at 37 °C and 200 rpm to OD₆₀₀ ≈ 0.8, then induced with 1 mM IPTG for 4 h at 30 °C. Cells were harvested by centrifugation, lysed by sonication in lysis buffer (50 mM sodium phosphate, 300 mM NaCl, 10 mM imidazole, 5 mM benzamidine, 1 mM PMSF, pH 7), and cleared by centrifugation at 12,000 g for 20 min at 4 °C. Supernatants were subjected to Ni–NTA agarose affinity purification according to the manufacturer’s (Qiagen) recommendations. Purified fractions were dialyzed sequentially against 50 mM phosphate buffers of pH 6.5, 6.0, and 5.5.

### Alkalinization and ROS assays

Extracellular alkalinization assays were performed in tomato (*S. peruvianum*) wild type and *syr1* mutant cell suspension cultures as described (Wang et al., 2024)(Li et al., 2025). Briefly, pH was recorded at 10 s intervals following elicitor treatment in 10 mL aliquots of cell suspension seven days after subculture. Custom-synthesized peptides or purified PS proteins were added from 200-fold concentrated stock solutions prepared in ddH₂O or 50 mM phosphate buffer (pH 5.5), respectively.

Reactive oxygen species (ROS) production was measured in 4 mm tomato leaf discs excised from 3-week-old seedlings as described (Li et al., 2024). Briefly, discs were incubated overnight in 200 μL ddH₂O in a 96-well plate at 24 °C. The next day, ddH₂O was replaced with 190 μL of reaction buffer containing 20 μM luminol (L-012, Wako) and 2 μg mL⁻¹ horseradish peroxidase. Elicitors were added from 20-fold concentrated stocks, and luminescence was recorded as relative light units (RLU) using a Tecan Spark® plate reader with emission detection between 415–530 nm.

### RNA-seq sample preparation and processing

Two-week-old tomato seedlings were sprayed with either ddH₂O (mock) or systemin peptides at 1 µM in aqueous solution. Leaves were harvested at 20 min, 1 h, 3 h, and 8 h post-treatment. RNA extraction and library construction followed established protocols (Yuan et al., 2023). Specifically, total RNA was isolated using a guanidine hydrochloride-based extraction buffer (8 M guanidine hydrochloride, 20 mM MES, 20 mM EDTA), 750 ng RNA per sample were barcoded, pooled, and sequenced across five single-end lanes on an Illumina HiSeq 3000, targeting ∼10 million reads per sample.

FASTQ files were merged and aligned to the SL4.0 tomato reference genome (https://doi.org/10.1101/767764) with ITAG4.0 annotation using RSEM with Bowtie2 (default parameters). On average, ∼7 billion reads per sample were mapped to the tomato genome, yielding an average mapping rate of 82.5%, and 34,075 gene models were detected across all samples. Libraries exceeding 8 million mapped reads were downsampled with seqtk. Transcripts mapping to organellar genomes, rDNA clusters, transposable elements, pseudogenes, or with effective lengths shorter than 150 nt were excluded. Transcript abundance was re-estimated in transcripts per million (TPM). Libraries with < 2 million mapped reads and those identified as outliers by principal component analysis of log₂(TPM) values were removed. Genes were retained if they had a trimmed mean log₂(TPM) > 0.3 and a coefficient of variance > 0.15.

### Differential expression analysis

Genes differentially expressed compared to the mock control (DEGs) were identified using the edgeR package (Robinson et al., 2010; McCarthy et al., 2012) in Bioconductor (https://bioconductor.org). Lowly expressed genes were filtered out, and library sizes were normalized using the Trimmed Mean of M-values (TMM) method. A negative binomial generalized log-linear model was fitted to estimate dispersion across biological replicates. The design included peptide-treated and mock plants at four time points (20 min, 1 h, 3 h, 8 h). Quasi-likelihood F-tests were applied, and genes with false discovery rate (FDR) < 0.05 and |log₂ fold change| ≥ 1 were considered significantly differentially expressed.

### Time-course transcriptome dynamics and clustering

To capture dynamic transcriptional changes, gene expression trajectories were modeled using linear mixed-model splines (LMMS) (Straube et al., 2015) in R v4.0 (R Core Team, 2022) with the full model as:

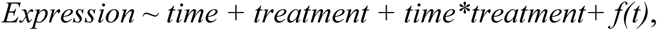

The full model included time, treatment, and their interaction, with transcript abundance fitted as a natural cubic B-spline function of time. Stepwise model selection was applied to prevent overfitting, and fitted expression values for each gene × treatment × time point were extracted for clustering.

Temporal expression patterns were grouped using K-means clustering. Each gene was represented by its LMMS-fitted values across all treatments and time points. The optimal number of clusters was determined by maximizing the Dunn index while constraining no more than 25% of clusters to contain fewer than 5% of genes. Clusters in which transcript trajectories did not co-vary with time under any treatment were discarded, and the remaining genes were re-clustered using the same criteria to obtain robust expression groups.

### Gene ontology enrichment analysis

Functional enrichment of differentially expressed genes was assessed using the PANTHER Classification System v14.0 (Overrepresentation Test). For each DEG set (per time point and direction of change), *Solanum lycopersicum* gene identifiers were tested against the background of all expressed genes retained after RNA-seq filtering. Enrichment was performed for GO Biological Process categories using Fisher’s exact test with Benjamini-Hochberg FDR correction. Terms with FDR < 0.05 were considered significant. Enriched GO terms were plotted and summarized in Figure 3D, Supplementary Figure 4C, and detailed in Supplementary Tables S1 and S2.

### Quantitative reverse transcriptase PCR (qRT-PCR)

Tomato seedlings were treated with peptides as described above, and leaves were harvested for RNA extraction. Total RNA was isolated using a guanidine hydrochloride-based extraction buffer followed by Column-based (EconoSpin^TM^, Epoch Life Science Inc., 19200-250) clean-up procedure. Extracted RNA was treated with DNase I (Thermo Fisher Scientific, EN0521) to remove genomic DNA contamination. 2 μg of RNA was reverse-transcribed using RevertAid reverse transcriptase (Thermo Fisher Scientific, EP0442) and oligo(dT) primers. Quantitative PCR was performed on a Bio-Rad CFX Connect real-time PCR instrument using SYBR Green (Invitrogen, S7563). Gene expression was quantified using the ΔΔCt method, with normalization to the expression levels of three housekeeping genes: *ACTIN2*, *UBIQUITIN10 (UBQ10)*, and *elongation factor 1α (EF1α)*.

### Phytohormone measurement

Tomato leaves were chopped into ∼2×2 mm leave pieces using a herb scissor and floated on ddH_2_O overnight. Ethylene emission was assayed by placing three leaf pieces in 6-ml tubes with 500 μl of ddH_2_O or ddH_2_O containing the appropriate concentration of elicitors. After tubes were securely sealed with rubber caps, 1 mL of air in the headspace was taken with a needled syringe at appropriate time points. Headspace samples were analyzed by gas chromatography (Shimadzu, GC-14A) for ethylene quantification as described (Wang et al., 2025).

Concentrations of JA JA-Ile and SA were quantified from 100 mg leaf tissue harvested and flash-frozen in liquid nitrogen one hour after elicitor or mock treatment. The sample was homogenized in 1 mL ethyl acetate spiked with deuterated phytohormones as internal standards (D6-JA, D6-JA-Ile, D4-SA) with 6-7 ceramic beads (Zirconox®) in a Bead Mill 24 (Fisherbrand^TM^, 2 x 20 sec at 4 m/s). After centrifugation, the pellet was re-extracted (15 sec at 4 m/s) in 1 ml ethyl acetate and centrifuged. The combined supernatants were dried in a vacuum concentrator (Eppendorf, Hamburg, Germany) and reconstituted in 400 µL 70% methanol (v/v) with 0.1% formic acid by vortexing for 10 min. The extracts were stored at −20°C until 4 µL were subjected to UPLC-ESI-MS/MS (Agilent 1290 II Infinity UHPLC coupled to a SCIEX QTRAP 6500+) analysis. Phytohormones were separated on a C18 column (Agilent ZORBAX Eclipse Plus, C18, 95 Å, 1,8 μm, 2.1 × 50 mm, 40°C) at 40°C with flow rate 0.35 mL/min with water and acetonitrile (both containing 0.2% formic acid) as solvents A and B in gradient mode (solvent B: 0 min: 15%; 2 min: 35%; 5.6 min: 38%; 6 min: 95%; 8 min: 95%; 8.2 min: 15%). Phytohormones were detected in negative ionization mode with parent/daughter ion selections of 209/59 (JA), 215/59 (D6-JA), 322/130 (JA-Ile), 328/130 (D6-JA-Ile), 263/153 (ABA), 269/159 (D6-ABA), 137/93 (SA), and 141/97 (D4SA), and quantified by peak area integration relative to the internal standards.

### Co-IP and Co-IP/MS

Transgenic SYR1-GFP plants (Wang *et al*., 2018) were pre-selected for comparable GFP fluorescence intensity. 100 nM peptide was vacuum-infiltrated into detached leaves in a desiccator at 100 mbar. Fully infiltrated leaves were collected immediately into 2 mL-Eppendorf tubes containing a metal bead, and frozen in liquid nitrogen. Frozen samples were homogenized in a bead beater (TissueLyser LT, Qiagen) and total protein was extracted in solubilization buffer (25 mM Tris, 150 mM NaCl, 1% NP40, 0.5% sodium deoxycholate, 2 mM DTT, pH 8.0) complemented with protease inhibitor mix P (SERVA). After 1 hour of gentle shaking at 4°C, protein extracts were incubated with 15 μl pre-equilibrated GFP-Trap^®^ Magnetic Agarose (ChromoTek). For MS analysis, samples were subjected to on-bead digestion with trypsin (0.5 μg/ μl, GE Life Sciences) in digestion buffer (6M Urea, 50mM Tris-HCl, pH 8.5). Tryptic peptides were analyzed by LC-MS/MS as described (Stührwohldt et al., 2020). Protein identification was carried out by MaxQuant version 1.5.3.8 against the tomato genome as reference (annotation version ITAG4.0).

### Confocal microscopy and FRAP analyses

Confocal imaging was performed using a Zeiss LSM700 microscope equipped with an LD LCI Plan-Apochromat 40×/1.2 water immersion objective and a 488 nm argon laser. For FRAP experiments, sixty consecutive images were acquired over ∼300 s. The region of interest (ROI) was photobleached with the 488 nm laser line at full power after the fourth scan, resulting in ≥80% reduction in fluorescence intensity. Recovery was quantified as the relative increase in fluorescence from the fifth scan (first post-bleach) to the final scan, normalized to the intensity difference between the last pre-bleach and the first post-bleach scan. Fluorescence images were processed and rendered with false color for visualization using ZEN Blue (v3.6). To minimize sample drift, 4 mm leaf discs were collected with a biopsy punch and mounted in water in imaging chambers.

### Insect feeding assays

Detached leaves from 1-month-old tomato plants (*Solanum lycopersicum* cv. UC82B) were pre-treated overnight with either ddH₂O (mock), 1 μM SYS1, or 1 μM SYS2 peptide solutions (Pepmic). Leaves were placed in feeding cages with their petioles submerged in the corresponding treatment solution to allow uptake via vascular transport. Sixty second-instar *Spodoptera exigua* larvae were used per treatment, distributed as 20 larvae per cage (three cages total). Feeding assays were conducted in controlled growth chambers (16 h photoperiod, 26 °C). Pre-treated leaves were replaced daily throughout the assay. Larval mass was measured every second day from day 4 to day 10 of feeding, and mortality was calculated as the proportion of dead larvae relative to the initial population size (n = 60) at each measurement point. Insect images were taken with a Nikon D5600 DSLR camera with 18–55 mm f/3.5–5.6 auto focus-P Nikkor Zoom lens.

## Acknowledgements

The work was supported by grants from the German Research Foundation (DFG) to ASt and ASc (CRC 1101/D06) and GF (CRC 1101/D05). The Sciex Qtrap 6500+ mass spectrometer was funded in part by the German Research Foundation (DFG-INST 36/174-1 FUGG). We thank Ursula Glück-Behrens and Bianca Bukowski for maintenance of the *S. peruvianum* cell cultures, and Jens Pfannstiel, Philipp Hubel, Iris Klaiber and Berit Würtz at the Core Facility Hohenheim for mass spectrometry analysis.

---

**Figure 1. Prosystemin proteins differ in length and amino acid composition.**

A. Phylogenetic analysis showing the relationship between PS1-4 protein sequences. The phylogenetic tree was built using the neighbor-joining algorithm by Clustal Omega. Numbers represent the estimated genetic divergence among PS protein sequences. **B**. Sequence alignment of tomato prosystemin proteins PS1-PS4 (Clustal Omega). Identical, highly and partially conserved residues are marked by asterisks (*), colons (:), and periods(.) respectively. Conserved SYS peptide regions are boxed in red. Length of the sequences (number of residues) is indicated. **C**. Transcripts levels of PS genes in leaves of wounded tomato plants. Data are the same as those used for Figure 1C, but shown as absolute number of transcripts (log2 of normalized transcripts per million (TPM)), rather than log2 fold change.

**Supplementary Figure 2. Genotype of CRISPR/Cas9-generated SYR receptor mutants**

Results are shown for five single and double mutants as indicated on the left. *SYR* loci were sequenced and chromatograms obtained for the mutants were compared to the SYR reference sequences on top. Guide RNA target sites are shown in brown type font, PAM sequences are boxed, Cas9 cleavage sites are marked. Two independent *syr1* single mutants were obtained (lines 6-27 and 3-100) which carry mutations at the target sites of both guide RNAs in the CRISPR construct, resulting in premature stop codons and truncated SYR1 proteins of 104 amino acids. The two *syr2* single mutants (lines 3-10 and 6-15) harbor mutations at one of the two gRNA target sites, producing truncated SYR2 proteins of 140 amino acids. The *syr1syr2* double mutant (line 2-2) contains large deletions in both the *SYR1* and *SYR2* loci, yielding truncated proteins of 286 and 59 amino acids, respectively.

**Supplementary Figure 3. SYS peptides elicit reactive oxygen species production in Nicotiana benthamiana.**

ROS production kinetics in SYR1-expressing *N. benthamiana* leaves following SYS peptide treatments. *35S:SYR1-GFP* was agroinfiltrated for transient expression in leaves 5 days prior to peptide elicitation.

**Supplementary Figure 4. SYS peptide-induced transcriptome reprogramming captured by RNA-seq.**

**A,B.** Experimental design. A schematic representation of peptide application to wild-type tomato seedlings and sampling time is shown in (**A**), the experimental workflow from sample collection to identification of differentially expressed genes in (**B**). **C.** Bubble plot showing Gene Ontology (GO) terms enriched among DEGs co-regulated by SYS1/2/3/4. Enrichment levels and statistical confidence (-lgFDR) are shown on the y- and x-axis, respectively. Bubble size reflects the number of genes associated with each GO term. **D.** Representative gene clusters displaying SYS2-specific transcriptional regulation. K indicates the cluster number; n indicates the number of genes the cluster contains.

**Supplementary Figure 5. Genes associated with ethylene synthesis and signaling are preferentially upregulated by SYS2, SYS3, and SYS4**

The heatmap shows SYS1/2/3/4 induced expression profiles for genes involved in ethylene biosynthesis, perception, and signaling. Gene number is indicated on the right; gene function is color-coded as transcription factors, biosynthetic enzymes, receptors, pathway regulators, or ethylene-responsive genes. Several genes are annotated as ethylene “overproducers” or pathway-related components. Red boxes highlight gene clusters preferentially induced by SYS2, SYS3, and SYS4. The dendrogram shows the hierarchical clustering result of the expression profiles of presented genes.

**Supplementary Figure 6. Systemin peptides do not induce salicylic acid accumulation.**

Salicylic acid (SA) levels measured 1 h after SYS1, SYS2, or mock treatment in wild-type, *syr1*, *syr2*, and *syr1syr2* mutant leaves of n = 8 plants. ns, not significant (P ≥ 0.05; two-tailed Student’s t-test). Equivalent results were obtained in three independent repetitions.

**Supplementary Figure 7. SYS1 -treated leaves are refractory to a second treatment with SYS1 but still respond to SYS2, SYS3 or SYS4.**

Ethylene production was measured in hourly intervals in tomato leaves that were all pretreated with SYS1 at t = - 4hrs, which then received a second treatment with SYS1 (red), SYS2 (blue), SYS3 (green), SYS 4 (yellow) or mock (gray) at t = 0. The experimental setup corresponds to Figure 4E.

**Supplementary Figure 8. SYS peptides differentially promote SYR1–SERK co-receptor interactions.**

Co-immunoprecipitation analysis of SYR1 interaction with SERK3A or SERK3B following SYS peptide treatments. SYR1-sfGFP was co-expressed with Myc-tagged SERK3A or SERK3B in *N. benthamiana* leaves by agro-infiltration. After five days, 100 nM peptide solutions were infiltrated as indicated. Total protein extracts were immunoprecipitated with GFP-trap beads, then detected with anti-GFP and anti-Myc on western blots. An anti-Myc blot and Coomassie (CBB)-stained gel of the input fractions are shown as loading controls. The assay was performed two times with equivalent results.

## Notes

### Competing Interest Statement

The authors have declared no competing interest.

