## Supplementary figures and tables for "Recruitment of SERK co-receptors determines signaling specificity within the systemin peptide family"

### Supplemental Figure 1

**A**

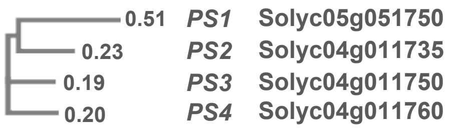

**B**

```
PS1      -----------MGTPSYDIKNKGDDMQEEPKVKLHHEKGGDEKE---KIIEKETPSQDIN   46
PS4      MANFFID-IKKIEDLFVYIVSCGKSSKEGEENDG--------------------------   33
PS2      MTDFITYTKKKIEDLSVYIVSCGKSYKEPARDVPEEDQAGPVTNITPIRFSKEPPR---N   57
PS3      MAEFIEDTKNKNEDPSISIVSDGKSSNEPSKVD---------------------------   33
                                  * . *.. :*   .

PS1      NKDTISSYVLRDDTQEIPKMEHEEGGYVKEKIVEKETISQYIIKIEGDDDAQEKLKVEYE   106
PS4      ------------------------------------------------------------   33
PS2      DQEPTSDHVLEDDFGIIKVFP-----------------------SGNAPPI------NN   87
PS3      -------Q-----------------------------------VVETQAE------NS   43

PS1      EEEYEKEKIVEKETPSQDINNKGDDAQEKPKVEHEEGDDKETPSQDIIKMEGEGALEITK   166
PS4      ------------------------------------------------------------   33
PS2      AQNQEEENIIAQQ--SQELDKSKGNSQEQKEIDD----------DEDF------------   123
PS3      TQNQEKENIIAQQ--GQEVDKSKENSQKQEQEQT----------EEE-------------   78

                      SYS
PS1      VVCEKIIVREDLAVQSKPPSKRDPPKMQTDNNKL       200
PS4      ----------GRANVSRPPGQRPAPRPQTGR---       54
PS2      KTLPKGFIPEERAKFSTPPGKRSAPRPKTG----       153
PS3      EEEEENANEEDRANFSTRPGKRSAPKPKTG----       108
                     *  *  *.:*  *: :*.
```

**C**

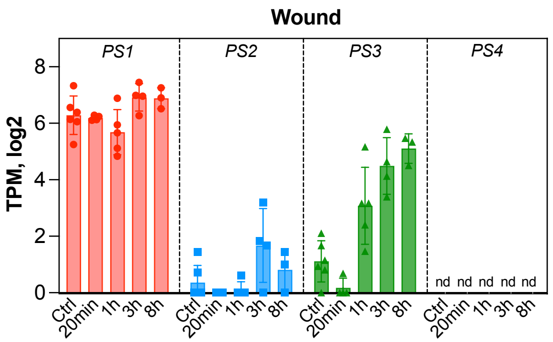

### Supplemental Figure 2

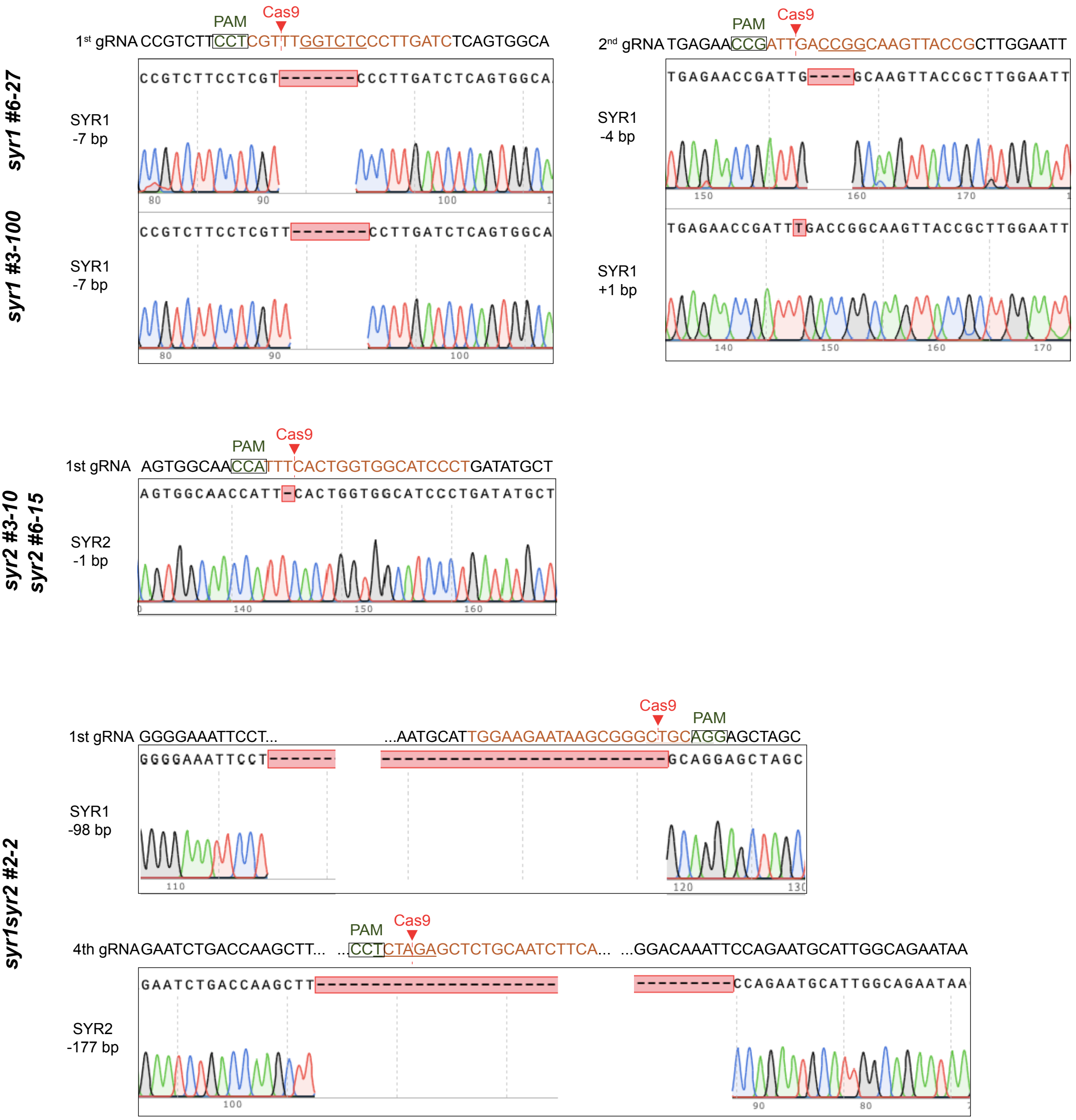

Supplemental Figure 3

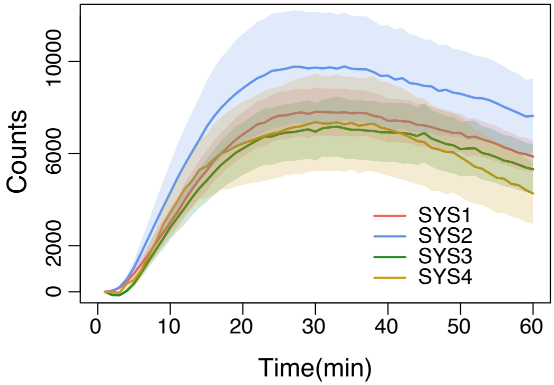

Supplemental Figure 4

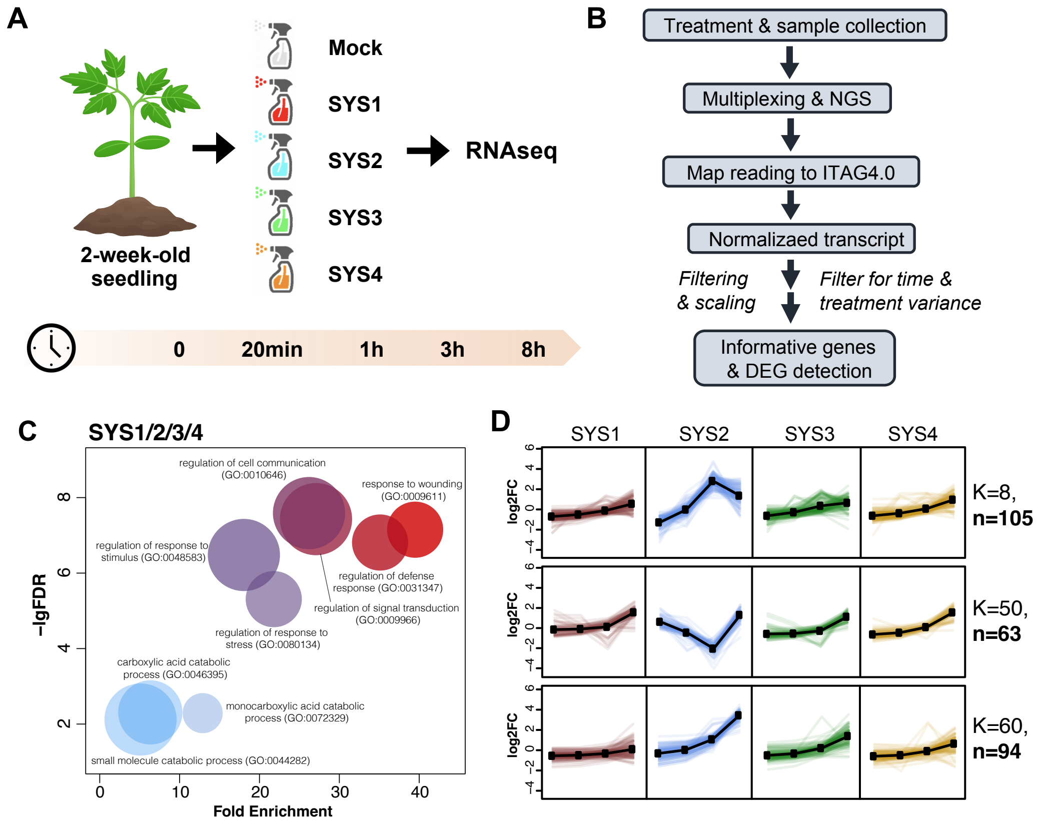

### Supplemental Figure 5

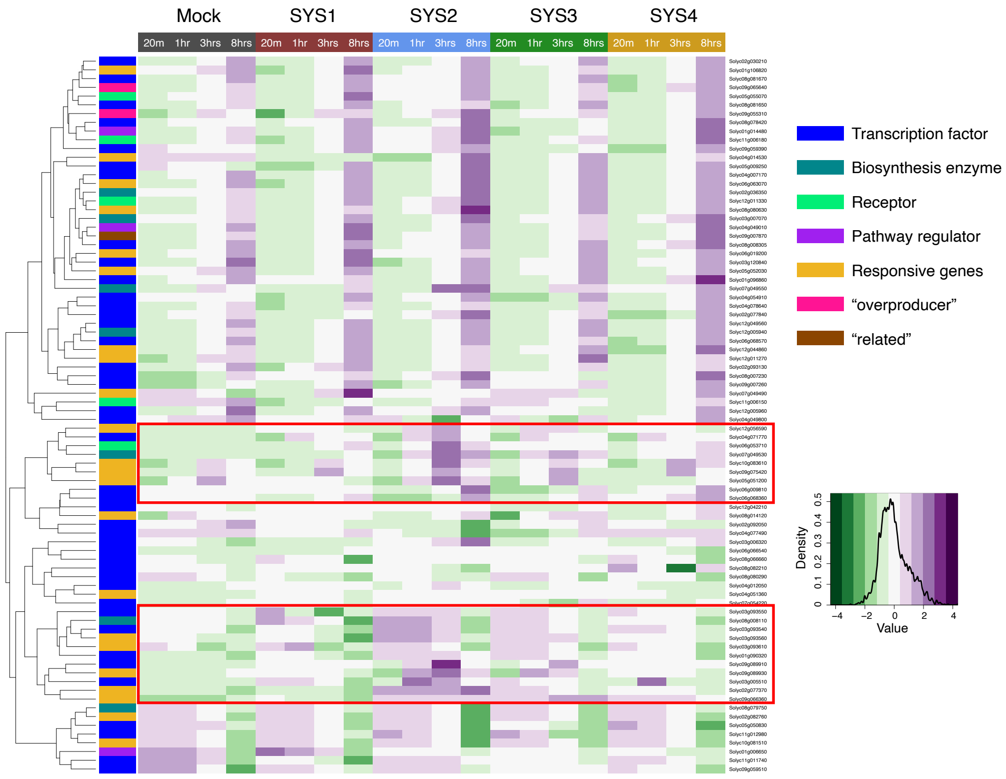

Supplemental Figure 6

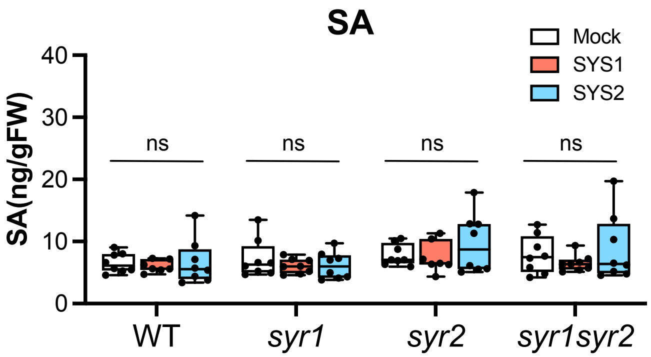

### Supplemental Figure 7

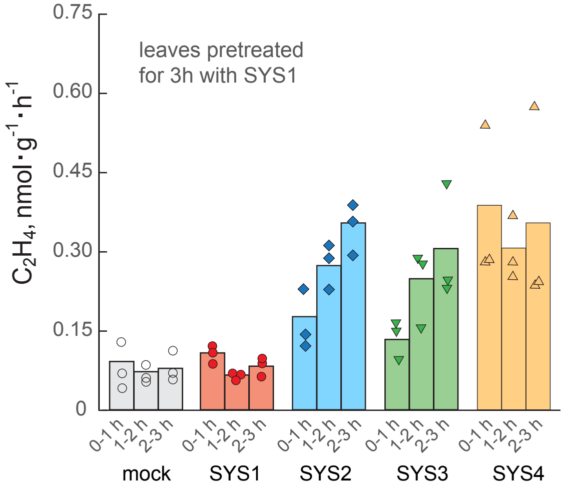

Supplemental Figure 8

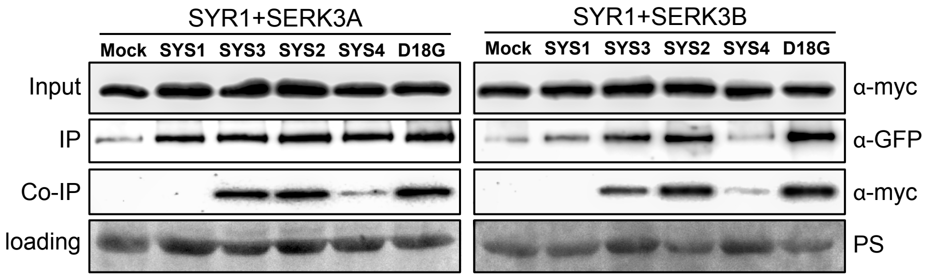

**Supplementary Table S1**

SYS1-specific DEGs at 3hr overrepresentation test by Panther GOslim

| **PANTHER GO-Slim Biological Process** | **fold Enrichment** | **raw P-value** | **FDR** |
|---|---|---|---|
| response to wounding (GO:0009611) | 38.9 | 9.93E-05 | 1.26E-02 |
| fatty acid beta-oxidation (GO:0006635) | 29.64 | 2.04E-04 | 1.83E-02 |
| regulation of defense response (GO:0031347) | 29.64 | 2.04E-04 | 1.72E-02 |
| fatty acid catabolic process (GO:0009062) | 25.93 | 2.91E-04 | 2.34E-02 |
| lipid oxidation (GO:0034440) | 19.76 | 7.12E-05 | 9.86E-03 |
| monocarboxylic acid catabolic process (GO:0072329) | 17.29 | 8.73E-04 | 4.92E-02 |
| regulation of signal transduction (GO:0009966) | 15.66 | 1.65E-04 | 1.80E-02 |
| regulation of cell communication (GO:0010646) | 15.37 | 1.77E-04 | 1.80E-02 |
| regulation of signaling (GO:0023051) | 15.37 | 1.77E-04 | 1.69E-02 |
| carboxylic acid catabolic process (GO:0046395) | 13.08 | 1.74E-06 | 5.29E-04 |
| organic acid catabolic process (GO:0016054) | 13.08 | 1.74E-06 | 4.41E-04 |
| cellular amino acid catabolic process (GO:0009063) | 12.2 | 4.08E-04 | 3.10E-02 |
| lipid modification (GO:0030258) | 10.92 | 6.08E-04 | 4.41E-02 |
| fatty acid metabolic process (GO:0006631) | 10.78 | 6.37E-04 | 4.41E-02 |
| regulation of response to stimulus (GO:0048583) | 10.64 | 6.68E-04 | 4.42E-02 |
| small molecule catabolic process (GO:0044282) | 10.12 | 1.96E-06 | 4.26E-04 |
| aromatic compound catabolic process (GO:0019439) | 7.2 | 7.84E-04 | 4.78E-02 |
| organic cyclic compound catabolic process (GO:1901361) | 7.06 | 8.58E-04 | 5.03E-02 |
| monocarboxylic acid metabolic process (GO:0032787) | 6.63 | 1.16E-04 | 1.36E-02 |
| cellular amino acid metabolic process (GO:0006520) | 6.59 | 3.96E-05 | 6.70E-03 |
| carboxylic acid metabolic process (GO:0019752) | 6.08 | 4.40E-08 | 6.70E-05 |
| organic acid metabolic process (GO:0006082) | 6.04 | 4.73E-08 | 3.61E-05 |
| oxoacid metabolic process (GO:0043436) | 6.04 | 4.73E-08 | 2.40E-05 |
| oxidation-reduction process (GO:0055114) | 4.82 | 7.51E-04 | 4.77E-02 |
| small molecule metabolic process (GO:0044281) | 4.08 | 1.34E-06 | 5.08E-04 |
| organic substance catabolic process (GO:1901575) | 3.64 | 2.13E-05 | 4.06E-03 |
| catabolic process (GO:0009056) | 3.38 | 4.91E-05 | 7.47E-03 |

**Supplementary Table S2**

SYS2-specific DEGs at 3hr overrepresentation test by Panther GOslim

| PANTHER GO-Slim Biological Process | fold Enrichment | raw P-value | FDR |
|---|---|---|---|
| programmed cell death (GO:0012501) | 21.24 | 3.01E-04 | 1.91E-02 |
| cell death (GO:0008219) | 21.24 | 3.01E-04 | 1.83E-02 |
| innate immune response (GO:0045087) | 19.11 | 4.25E-04 | 1.98E-02 |
| immune system process (GO:0002376) | 19.11 | 4.25E-04 | 1.92E-02 |
| immune response (GO:0006955) | 19.11 | 4.25E-04 | 1.86E-02 |
| post-translational protein targeting to endoplasmic reticulum membrane (GO:0006620) | 14.7 | 9.78E-04 | 3.18E-02 |
| tricarboxylic acid cycle (GO:0006099) | 11.08 | 4.19E-04 | 2.02E-02 |
| protein targeting to ER (GO:0045047) | 9.44 | 7.91E-04 | 2.83E-02 |
| establishment of protein localization to endoplasmic reticulum (GO:0072599) | 9.44 | 7.91E-04 | 2.76E-02 |
| clathrin-dependent endocytosis (GO:0072583) | 9.1 | 9.11E-04 | 3.03E-02 |
| amino acid transmembrane transport (GO:0003333) | 8.74 | 1.44E-05 | 1.55E-03 |
| organic acid transmembrane transport (GO:1903825) | 7.96 | 2.69E-05 | 2.51E-03 |
| carboxylic acid transmembrane transport (GO:1905039) | 7.96 | 2.69E-05 | 2.35E-03 |
| receptor-mediated endocytosis (GO:0006898) | 7.96 | 1.52E-03 | 4.35E-02 |
| amino acid transport (GO:0006865) | 7.19 | 5.26E-05 | 3.87E-03 |
| organic anion transport (GO:0015711) | 6.91 | 6.35E-06 | 8.07E-04 |
| protein transmembrane transport (GO:0071806) | 6.13 | 1.32E-03 | 3.94E-02 |
| intracellular protein transmembrane transport (GO:0065002) | 6.13 | 1.32E-03 | 3.86E-02 |
| carboxylic acid transport (GO:0046942) | 6.11 | 1.50E-04 | 1.05E-02 |
| organic acid transport (GO:0015849) | 6.11 | 1.50E-04 | 1.00E-02 |
| peptidyl-serine phosphorylation (GO:0018105) | 4.69 | 7.59E-04 | 2.87E-02 |
| peptidyl-serine modification (GO:0018209) | 4.69 | 7.59E-04 | 2.79E-02 |
| protein targeting (GO:0006605) | 4.14 | 7.45E-04 | 2.89E-02 |
| establishment of protein localization to organelle (GO:0072594) | 3.8 | 6.67E-04 | 2.66E-02 |
| nitrogen compound transport (GO:0071705) | 3.75 | 2.18E-08 | 3.05E-05 |
| organic substance transport (GO:0071702) | 3.37 | 5.50E-08 | 3.85E-05 |
| protein phosphorylation (GO:0006468) | 3.33 | 1.81E-05 | 1.81E-03 |
| transmembrane transport (GO:0055085) | 3.19 | 1.08E-05 | 1.26E-03 |
| phosphorylation (GO:0016310) | 3.09 | 4.61E-05 | 3.58E-03 |
| intracellular protein transport (GO:0006886) | 2.85 | 1.19E-03 | 3.60E-02 |
| protein transport (GO:0015031) | 2.81 | 8.71E-04 | 2.97E-02 |
| macromolecule localization (GO:0033036) | 2.78 | 4.58E-05 | 3.76E-03 |
| establishment of protein localization (GO:0045184) | 2.73 | 1.11E-03 | 3.45E-02 |
| cellular macromolecule localization (GO:0070727) | 2.67 | 4.07E-04 | 2.11E-02 |
| protein localization (GO:0008104) | 2.67 | 4.07E-04 | 2.03E-02 |
| signal transduction (GO:0007165) | 2.59 | 3.81E-04 | 2.13E-02 |
| signaling (GO:0023052) | 2.58 | 3.91E-04 | 2.10E-02 |

**Supplementary Table S2 (continued)**

SYS2-specific DEGs at 3hr overrepresentation test by Panther GOslim

| | | | |
|---|---|---|---|
| phosphate-containing compound metabolic process (GO:0006796) | 2.58 | 1.97E-06 | 3.06E-04 |
| phosphorus metabolic process (GO:0006793) | 2.53 | 2.93E-06 | 4.09E-04 |
| cell communication (GO:0007154) | 2.47 | 6.43E-04 | 2.64E-02 |

**Supplementary Table S3**

List of primers used in this study

| Gene ID | Primer Name | Sequence(5' to 3') | Purpose |
|---|---|---|---|
| Solyc04g011750 | FLAG_SYS3_Fw | TACAAGGATGATGACGACAAAGCTGAGTTCATTGAAGATACA | Create FLAG, sfGFP fusion |
| | SYS3_linker_Rv | TTGCGCGCCCGCGGTGGCTGCCGCACCAGTCTTAGGTTTAGGAG | |
| Solyc04g011735 | FLAG_SYS2_Fw | ACTACAAGGATGATGACGACAAAACTGATTTCATCACATATACA | Create FLAG, sfGFP fusion |
| | SYS2_linker_Rv | GCGCCCGCGGTGGCTGCCGCACCAGTCTTAGGTCTAGGAGCT | |
| Solyc04g011760 | FLAG_SYS4_Fw | ACTACAAGGATGATGACGACAAAGCTAATTTCTTCATAGATATA | Create FLAG, sfGFP fusion |
| | SYS4_linker_Rv | GCGCCCGCGGTGGCTGCCGCTCGACCAGTCTGAGGCCTAGGA | |
| Solyc04g011750 | Fw_SYS3_XbaI | CCCTCTAGAATGGCTGAGTTCATTGAAGATACAAA | Add His tag |
| | Rv_SYS3_HindIII | CGCAAGCTTACCAGTCTTAGGTTTAGGAGCTGAT | |
| Solyc04g011735 | Fw_SYS2_XbaI | CCCTCTAGAATGACTGATTTCATCACATATACAAA | Add His tag |
| | Rv_SYS2_HindIII | CGCAAGCTTACCAGTCTTAGGTCTAGGAGCTGA | |
| Solyc04g011760 | Fw_SYS4_XbaI | CCCTCTAGAATGGCTAATTTCTTCATAGATATAAA | Add His tag |
| | Rv_SYS4_HindIII | CGCAAGCTTTCGACCAGTCTGAGGCCTAGGA | |
| Solyc03g082450 | DT824501-BsF | ATATATGGTCTCGATTgAGGGATGCCACCAGTGAAAGTT | Generate SYR2 single mutant |
| | DT824501-F0 | TgAGGGATGCCACCAGTGAAAGTTTTAGAGCTAGAAATAGC | |
| | DT824502-R0 | AACACGAGCTGACAGGACCTCTcAATCTCTTAGTCGACTCTAC | |
| | DT824502-BsR | ATTATTGGTCTCGAAACACGAGCTGACAGGACCTCTcAA | |
| Solyc03g082470 | DT824701-BsF | ATATATGGTCTCGATTgCGGTAACTTGCCGGTCAATGTT | Generate SYR1 single mutant |
| | DT824701-F0 | TgCGGTAACTTGCCGGTCAATGTTTTAGAGCTAGAAATAGC | |
| | DT824702-R0 | AACCGTTTGGTCTCCCTTGATCcAATCTCTTAGTCGACTCTAC | |
| | DT824702-BsR | ATTATTGGTCTCGAAACCGTTTGGTCTCCCTTGATCcAA | |
| Solyc03g082450/ Solyc03g082470 | 1222-Sl03g082470T1-BsF1 | ATATATGGTCTCGATTgGGAAGAATAAGCGGGCTGCGTT | Generate SYR1,SYR2 double mutant |
| | 1222-Sl03g082470T1-F0 | TgGGAAGAATAAGCGGGCTGCGTTTTAGAGCTAGAAATAGC | |
| | 1960-Sl03g082470T2-BsF2 | ATATTATTGGTCTCAAGATTgATGCTTGTAAAGCTGCAGCGTT | |
| | 1960-Sl03g082470T2-F0 | TgATGCTTGTAAAGCTGCAGCGTTTTAGAGCTAGAAATAGC | |
| | 337-Sl03g082450T3-BsF3 | ATATTATTGGTCTCAGTGATTgGTGTCAAGTTGGCTGCAGTGTT | |
| | 337-Sl03g082450T3-F0 | TgGTGTCAAGTTGGCTGCAGTGTTTTAGAGCTAGAAATAGC | |
| | 1113-Sl03g082450T4-R0 | AACCTAGAGCTCTGCAATCTTCcAATCACTACTTCGACTCTAGCTGTAT | |
| | 1113-Sl03g082450T4-BsR | ATTATTGGTCTCTAAACCTAGAGCTCTGCAATCTTCc | |

| | | | |
|---|---|---|---|
| Solyc03g082450/ Solyc03g082470 | 82450gRNAF | ATTGAGGGATGCCACCAGTGAAAG | CRISPR construct check |
| | U6-29p-R | AGCCCTCTTCTTTCGATCCATCAAC | |
| | 82470gRNAF | ATTGCGGTAACTTGCCGGTCAATG | |
| | U6-1t-F | GCTAAGACAAAGTGATTGGTCCGTT | |
| | 1113-Sl03g082450T4-BsR | ATTATTGGTCTCTAAACCTAGAGCTCTGCAATCTTCc | |
| Solyc03g082450 | 82450CRISPRF-136 | TCTGTCTCGCACTGCCAATGG | Genotyping SYR2 |
| | 82450CRISPRR-737 | CCAAGATGAGCAGATGATGCAGAG | |
| Solyc03g082470 | 82470CRISPRF1-172 | AAGGGTGTTACCTGCTATTCAGAC | Genotyping SYR1 |
| | 82470CRISPRR1-456 | GTAACCCAACTCAAGGTATACAAGC | |
| | 82470CRISPRF2-1071 | CACTTCTCTTGCTGAGATAAGTCTCG | |
| | 82470CRISPRR2-1427 | AGTTTAGAAGGAATCTGTCCACTGAAG | |
| Solyc03g082450/ Solyc03g082470 | syr1syr2_1st gRNA F | CAAGTCAGTTGGACGTCTGAAGAAT | Genotyping SYR1SYR2 |
| | syr1syr2_2nd gRNA R | AATTTGTCAAGATTGCCTAGGCACC | |
| | syr1syr2_3rd gRNA F | CCTGCTATTCAGACACAACTTGTCAT | |
| | syr1syr2_4th gRNA R | GCCAAAGAGAGTAACACAAGTTTCGT | |
| Solyc02g082830 | 02g082830_Fw | ACCCGCCATAACTACCAAAC | qRT-PCR |
| | 02g082830_Rv | GGCTGAGCAGGAAATAGACATAG | |
| Solyc04g040180 | 04g040180_Fw | TTTGGGATGTTGGTACTGGTAG | qRT-PCR |
| | 04g040180_Rv | CAAGCTGCTTTGGACTTGTG | |
| Solyc07g047800 | 07g047800_Fw | CTTTGTCAGTGGGTAAGGAGAG | qRT-PCR |
| | 07g047800_Rv | CAGTGGAAGCCAGATCCTTT | |
| Solyc07g049530 | 07g049530_Fw2 | CCATGTCCTAAGCCCGATTT | qRT-PCR |
| | 07g049530_Rv2 | GGCCACTCACTTTGTCATCT | |
| Solyc09g075820 | 09g075820_Fw | TGCCAATCTCGTCAACTACG | qRT-PCR |
| | 09g075820_Rv | GCGCCCAAAGTCAACAATAC | |
| Solyc06g005060 | elf1a 1272 | AGCCCATGGTTGTTGAGACCTTTG | qRT-PCR |
| | elf1a 1461R | TTCGAAACACCAGCATCACACTGC | |
| Solyc07g064130 | Ubi3 230F | CTCTTGCCGACTACAACATCC | qRT-PCR |
| | Ubi3 451R | AGCACCGCACTCAGCATTA | |
| Solyc03g078400 | SlACT 816F | CACTACTGCTGAACGGGAAAT | qRT-PCR |
| | SlACT 951R | CTGTCCATCTGGCAACTCATAG | |
| Solyc11g021060 | SlPI2 161 | GGATATGCCCACGTTCAGAAGGAA | qRT-PCR |
| | SlPI2 418R | AATAGCAACCCTTGTACCCTGTGC | |

**Supplementary Table S4**

Predicted template modeling (pTM) scores and interface predicted template modeling (ipTM) scores of the top 5 models generated by Alphafold3 (https://alphafoldserver.com/) for the ternary complexes comprising SYR1, SERK3A or 3B, and SYS1 or SYS1$^{GtoD}$.

| SYR1 complexed with | score | Model 0 | Model 1 | Model 2 | Model 3 | Model 4 |
|---|---|---|---|---|---|---|
| SERK3A-SYS1 | pTM<br>ipTM | 0.76<br>0.31 | 0.75<br>0.30 | 0.74<br>0.30 | 0.71<br>0.36 | 0.73<br>0.23 |
| SERK3A-SYS1$^{DtoG}$ | pTM<br>ipTM | 0.78<br>0.43 | 0.78<br>0.36 | 0.77<br>0.34 | 0.76<br>0.33 | 0.76<br>0.32 |
| SERK3A-SYS2 | pTM<br>ipTM | 0.87<br>0.83 | 0.87<br>0.83 | 0.87<br>0.84 | 0.87<br>0.83 | 0.86<br>0.83 |
| SERK3A-SYS2$^{GtoD}$ | pTM<br>ipTM | 0.75<br>0.56 | 0.73<br>0.56 | 0.74<br>0.55 | 0.73<br>0.55 | 0.72<br>0.54 |
| SERK3B-SYS1 | pTM<br>ipTM | 0.86<br>0.80 | 0.86<br>0.86 | 0.86<br>0.79 | 0.86<br>0.79 | 0.84<br>0.79 |
| SERK3B-SYS1$^{DtoG}$ | pTM<br>ipTM | 0.88<br>0.85 | 0.88<br>0.85 | 0.88<br>0.85 | 0.87<br>0.84 | 0.87<br>0.84 |
| SERK3B-SYS2 | pTM<br>ipTM | 0.87<br>0.83 | 0.87<br>0.82 | 0.87<br>0.82 | 0.86<br>0.82 | 0.87<br>0.82 |
| SERK3B-SYS2$^{GtoD}$ | pTM<br>ipTM | 0.87<br>0.82 | 0.86<br>0.81 | 0.86<br>0.81 | 0.86<br>0.81 | 0.86<br>0.81 |
| SERK1-SYS1 | pTM<br>ipTM | 0.80<br>0.55 | 0.79<br>0.53 | 0.73<br>0.27 | 0.71<br>0.24 | 0.72<br>0.24 |
| SERK1-SYS1$^{DtoG}$ | pTM<br>ipTM | 0.72<br>0.25 | 0.72<br>0.23 | 0.71<br>0.21 | 0.72<br>0.21 | 0.71<br>0.20 |
| SERK1-SYS2 | pTM<br>ipTM | 0.74<br>0.53 | 0.74<br>0.52 | 0.73<br>0.52 | 0.74<br>0.51 | 0.72<br>0.51 |
| SERK1-SYS2$^{GtoD}$ | pTM<br>ipTM | 0.72<br>0.54 | 0.72<br>0.53 | 0.72<br>0.51 | 0.71<br>0.51 | 0.71<br>0.49 |
